# Using a Whole Genome Co-expression Network to Inform the Functional Characterisation of Predicted Genomic Elements from *Mycobacterium tuberculosis* Transcriptomic Data

**DOI:** 10.1101/2022.06.22.497203

**Authors:** Jennifer Stiens, Yen Yi Tan, Rosanna Joyce, Kristine B. Arnvig, Sharon L. Kendall, Irene Nobeli

## Abstract

A whole genome co-expression network was created using *Mycobacterium tuberculosis* transcriptomic data from publicly available RNA-sequencing experiments covering a wide variety of experimental conditions. The network includes expressed regions with no formal annotation, including putative short RNAs and untranslated regions of expressed transcripts, along with the protein-coding genes. These unannotated expressed transcripts were among the best-connected members of the module sub-networks, making up more than half of the ‘hub’ elements in modules that include protein-coding genes known to be part of regulatory systems involved in stress response and host adaptation. This dataset provides a valuable resource for investigating the role of non-coding RNA, and conserved hypothetical proteins, in transcriptomic remodelling. Based on their connections to genes with known functional groupings and correlations with replicated host conditions, predicted expressed transcripts can be screened as suitable candidates for further experimental validation.

## INTRODUCTION

Tuberculosis continues to be a leading cause of death worldwide, causing over 1.5 million deaths, and infecting over 10 million people in 2020 (World Health Organization, 2021). The human-adapted pathogen causing tuberculosis, *Mycobacterium tuberculosis* (Mtb), has a complex lifestyle that requires rapid adaptation to host defences and immune pressure, including nutritional immunity, hypoxia and lipid-rich environments. In order to eradicate the disease, it is crucial to understand how the pathogen survives attacks from host immune cells and persists in an extended latent state inside the host. To adapt to these environmental challenges, bacterial cells must make complex transcriptomic adjustments, and these are thought to be complemented and fine-tuned by post-transcriptional regulation.

The mycobacterial genome produces a range of conditionally expressed transcripts, including non-coding RNA, short, unannotated ORFs and untranslated regions at the 5’ and 3’ end of protein-coding sequences, many of which are poorly annotated and understood. In this paper, we extend our focus to include ‘non-coding’ RNA (ncRNA), here referring to non-ribosomal RNA transcripts not known to be translated into peptides, such as short RNAs (sRNAs) acting on either distant or antisense mRNA targets and the expressed untranslated regions (UTRs) flanking coding regions (which may also contain short open reading frames (sORFs) upstream from coding regions). Non-coding RNA can alter the abundance of RNA and proteins by controlling mRNA stability, processing and access to ribosome binding sites. Discovering the contribution of the non-coding genome to specific adaptation-response pathways may improve our ability to design therapeutics and prevent the evolution of persistent phenotypes.

### Uncovering the role of non-coding RNA in adaptation and transcriptomic remodelling

The proportion of non-ribosomal, ncRNA in the Mtb transcriptome has been shown to increase in stationary and hypoxic conditions, indicating a potential role in adjusting to environmental cues (Aguilar-Ayala et al., 2017; K. B. Arnvig et al., 2011; Gerrick et al., 2018; Ignatov et al., 2015). Several mycobacterial ncRNA transcripts (particularly, sRNA) have been extensively studied and found to be associated with regulatory systems controlling adaptation to stress conditions or growth phase, linked to virulence pathways and access to lipid media (Arnvig et al., 2011; Gerrick et al., 2018; Girardin & McDonough, 2020; Mai et al., 2019; Moores et al., 2017; Solans et al., 2014). Non-coding regulation in Mtb appears to function quite differently compared to model organisms, eschewing the use of any known chaperone proteins for RNA-RNA interactions and with few sRNA homologs found outside the phyla (Gerrick et al., 2018; Mai et al., 2019; Schwenk & Arnvig, 2018). The discovery of ncRNA in Mtb has progressed using both molecular biology methods and high-throughput sequence-based approaches (reviewed in Schwenk & Arnvig, 2018) but uncovering the regulation and actions of a particular ncRNA is experimentally expensive and very few have been fully-characterised. Annotation of identified transcripts remains incomplete, as well, with only 30 listed in the Mtb H37Rv reference sequence (GenBank AL123456.3). Efforts to compile a comprehensive list of annotated ncRNAs for Mtb are impeded by non-standardised nomenclature, different standards of experimental validation, incomplete reference annotations (especially for the closely-related animal-adapted species of the Mycobacterium tuberculosis complex (MTBC)) and the variable expression of non-coding transcripts in response to different experimental conditions (Stiens et al., 2022).

Using RNA-sequencing (RNA-seq) data to predict ncRNA in the compact Mtb genome is challenging. Paradoxically, more sensitive, high-depth sequencing can make it more difficult to identify the small, low-abundance, functional transcripts above stochastic gene expression and technical noise. Parameters of detection must therefore be carefully considered for each dataset to account for variation in expression levels. Though RNA-seq-based ncRNA prediction algorithms are often assumed to overpredict putative ncRNAs, especially at the 5’ and 3’ ends of coding genes, there are biological and technical reasons for detecting abundant signal in the unannotated regions of the genome. Ribosome profiling (Ribo-seq) methods that sequence the ribosome-protected fragments of mRNA have identified actively translated RNA in the 5’ UTRs of annotated protein-coding mRNA transcripts (Canestrari et al., 2020; D’Halluin et al., 2022; Sawyer et al., 2021; Shell et al., 2015; Smith et al., 2022). These unannotated sORFs may represent functional peptides or function to regulate the translation of the downstream transcript; however, it is impossible to tell the difference between a putative ncRNA and a sORF from RNA-seq signal alone. Additionally, the 3’ ends in mycobacterial RNA-seq data often lack clear signal termination (Dar et al., 2016; D’Halluin et al., 2022; Lejars et al., 2019) and processing of transcripts at the 3’ end may be the norm (Wang et al., 2019). Finally, polycistronic transcripts often include non-coding sequence between the genes of an operon, and this may contain functional elements and/or processing sites (Martini et al., 2019).

The location of a transcription start site (TSS) in the 5’ end of a predicted transcript supports the biological relevance of a predicted ncRNA. However, the available lists of Mtb TSS sites (Cortes et al., 2013; Shell et al., 2015) have so far been mapped only in starvation and exponential growth and may not include TSSs that are expressed under different experimental conditions. New TSS maps, published subsequent to this analysis may increase the number of predicted transcripts with a TSS (D’Halluin et al., 2022). Furthermore, functional ncRNA elements generated from the 3’ UTRs of coding genes through RNase processing would presumably lack a TSS. 3’ UTRs that are functionally independent from their cognate coding sequence (CDS) have been identified in other bacteria (Desgranges et al., 2021; Menendez-Gil et al., 2020; Ponath et al., 2022). Therefore, it is important to consider predicted UTRs as separate annotated elements from protein-coding transcripts when quantifying differential expression.

To include a complete picture of the interaction of the non-coding genome with coding genes involved in adaptation pathways, we have generated a novel set of ncRNA sequence-based predictions (sRNAs and UTRs) from publicly available datasets using our in-house software package, *baerhunter* (Ozuna et al., 2019). Some of these predicted non-coding transcripts overlap with those of previous studies, but many represent novel predictions. The expression of these transcripts is quantified along with the protein-coding genes and used in network analysis to provide a more complete picture of the functional groupings involved in adaptation to environmental changes. Including a variety of culture conditions that replicate aspects of the host environment improves the chances that the expression of any ncRNA that is restricted to one or more conditions is included in the network (Ami et al., 2020).

### Using WGCNA to implicate functional associations of non-coding RNA

Weighted gene co-expression network analysis (WGCNA) (B. Zhang & Horvath, 2005) has been widely used to identify functional groups of genes, called ‘modules’, through the application of hierarchical clustering to differential expression levels of RNA transcripts in microarray or RNA-seq experiments. Recent studies have focussed entirely on the protein-coding portion of the transcriptome, using WGCNA with RNA-seq to cluster the differentially expressed genes of *Mycobacterium marinum* in response to resuscitation after hypoxia (Jiang et al., 2020) and *Mycobacterium aurum* infected macrophages (Lu et al., 2021). Mtb microarray data have been used to cluster protein-coding genes that show differential expression among clinical isoloates (Puniya et al., 2013) and in response to two different hypoxic models to identify potential transcription factors (Jiang et al., 2016). Another recent network analysis, using a matrix deconvolution method followed by module clustering, uses a large number of RNA-seq samples including deletion mutants, infection models and antibiotic-treated samples as well as restricted media and culture conditions (Yoo, et al., 2022). Here the authors identify 80 modules of protein-coding genes that each approximate an isolated source of variance, together estimated to account for 61% of the total variance seen in in the dataset. This proportion is reportedly lower than results from similar analyses in other organisms, potentially due to the bias in the types of conditions available in the database and/or the complex nature of regulation in Mtb (Yoo, et al., 2022). However, the contribution of regulatory ncRNA elements may be a considerable unexplored source of variance in this complex system. Here we use an alternative, complementary approach by including ncRNA, as well as annotated protein-coding genes, in the modules.

In this study, WGCNA was applied to multiple Mtb H37Rv datasets covering 15 different culture conditions replicating various growth conditions, nutrient sources and stressors encountered in the host environment. We present a global view of the non-coding genome across an extensive WGCNA network and interrogate selected modules to identify functional groupings between protein-coding and non-coding transcripts, as well as between well-characterised genes and those with little functional annotation. The correlation of the modules with the various conditions can identify participants in large-scale transcriptomic remodelling programs in response to changes in environmental conditions.

## MATERIALS AND METHODS

The overall workflow for this analysis is presented in Figure 1. All scripts for *baerhunter*, WGCNA and subsequent analysis are available at: https://doi.org/10.5281/zenodo.7319853.

**Figure 1.**
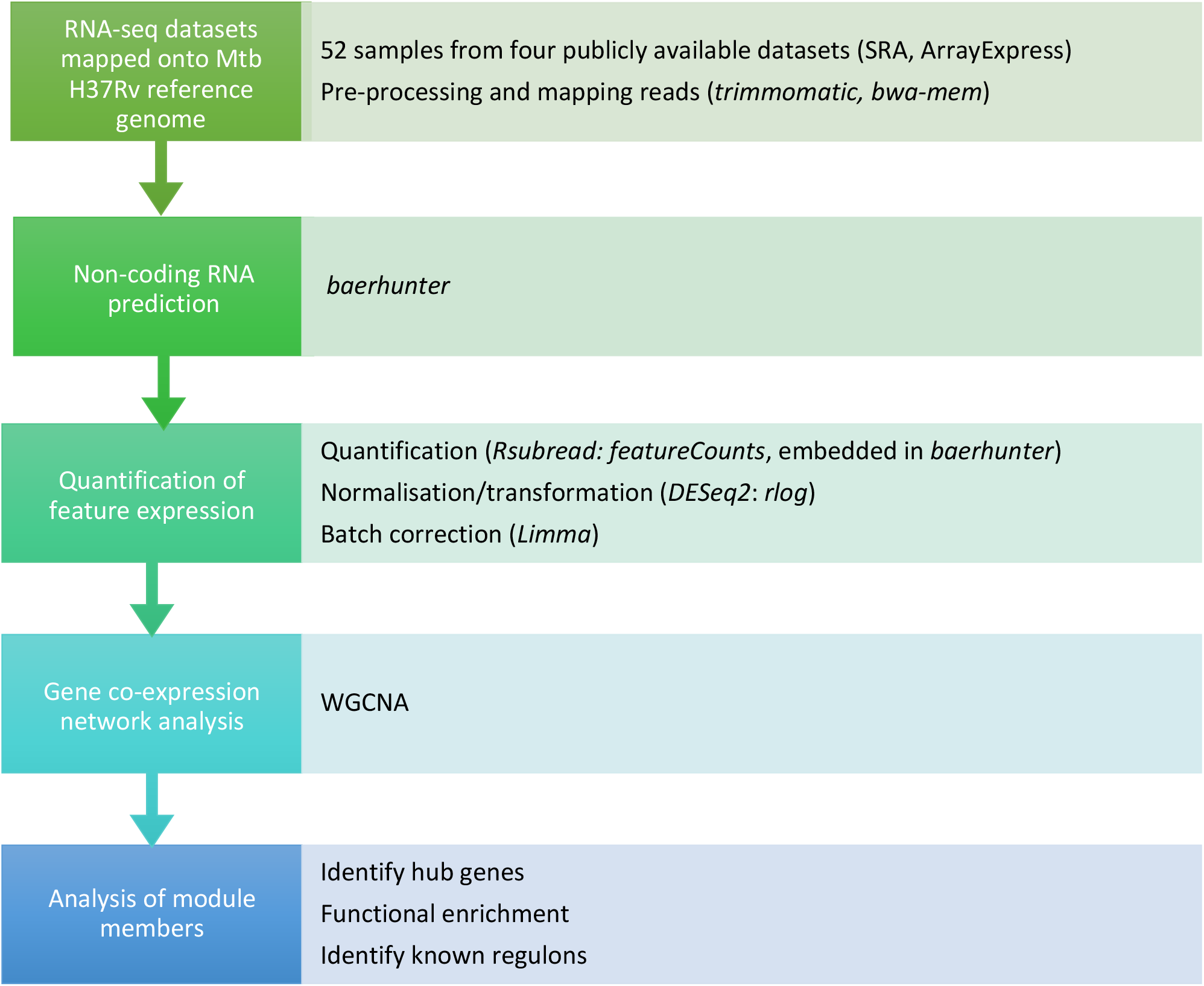
Analysis workflow

### Data Acquisition and Mapping

Datasets were downloaded from SRA (https://www.ncbi.nlm.nih.gov/sra/docs/) or Array Express (https://www.ebi.ac.uk/arrayexpress/) using the accession numbers listed in Table 1. To minimise batch effects and ensure compatibility with RNA prediction software, we limited analysis to datasets with similar library strategies. Samples were included based on inspection to confirm that 1) samples were from monocultures of wildtype Mtb H37Rv strain and 2) sequencing was using a paired-end, stranded protocol. Reads from samples that passed quality control thresholds were trimmed using *Trimmomatic* (Bolger et al, 2014) to remove adapters and low-quality bases from the 5’ and 3’ ends of the sequences. Trimmed reads were mapped to the H37Rv reference genome (GenBank AL123456.3) using *BWA-mem* in paired-end mode (Li, Heng, 2013). All samples had >70% percent reads mapped with an overall mean of ~ 27.75M mapped reads and a range of 3.97M to 60.68M mapped reads per sample (Supp Table 1, ‘Samples’ tab).

**Table 1.**
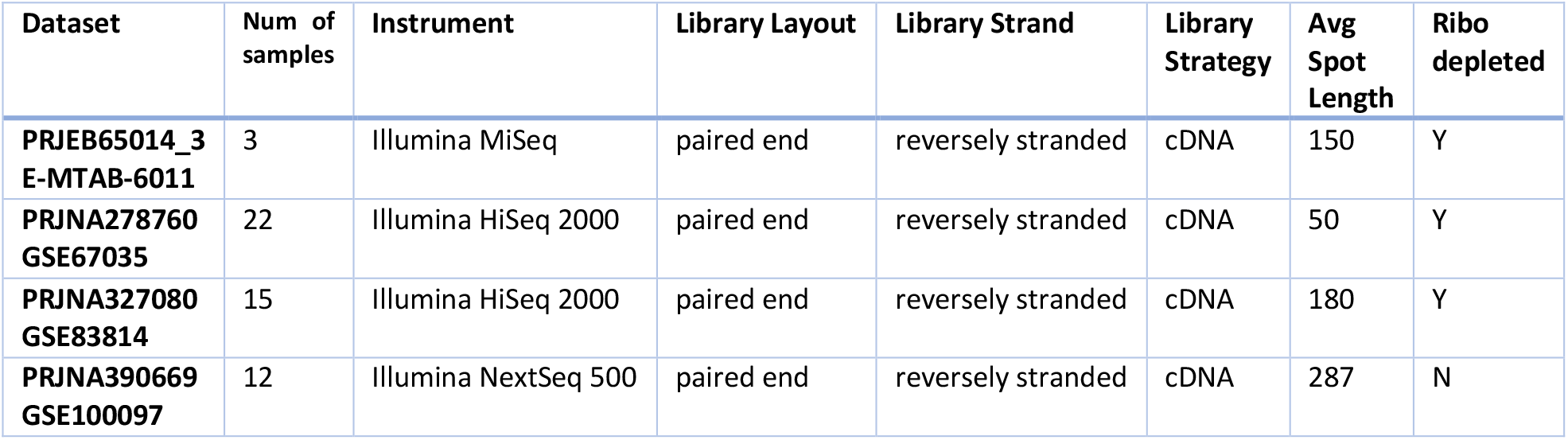
Datasets used in analysis. Accession numbers from SRA and Array Express.

| Dataset | Num of samples | Instrument | Library Layout | Library Strand | Library Strategy | Avg Spot Length | Ribo depleted |
| --- | --- | --- | --- | --- | --- | --- | --- |
| PRJEB65014_3 E-MTAB-6011 | 3 | Illumina MiSeq | paired end | reversely stranded | cDNA | 150 | Y |
| PRJNA278760 GSE67035 | 22 | Illumina HiSeq 2000 | paired end | reversely stranded | cDNA | 50 | Y |
| PRJNA327080 GSE83814 | 15 | Illumina HiSeq 2000 | paired end | reversely stranded | cDNA | 180 | Y |
| PRJNA390669 GSE100097 | 12 | Illumina NextSeq 500 | paired end | reversely stranded | cDNA | 287 | N |

### Non-coding RNA prediction

Each dataset was run through the R-package, *baerhunter* (Ozuna et al., 2019), using the *‘feature_file_editor’* function optimised to the most appropriate parameters for the sequencing depth (**https://doi.org/10.5281/zenodo.7319853**). ‘*Count_features’* and ‘*tpm_norm_flagging’* functions were used for transcript quantification and to identify low expression hits (less than or equal to 10 transcripts per million) in each dataset, which were subsequently eliminated. When viewed on a genome browser, coverage at the 3’ ends of putative sRNA and UTRs often appears to decrease gradually, with the actual end of the transcript appearing indistinct, compared to the 5’ end. Prokaryotic ncRNA transcripts may not demonstrate a clear fall-off of expression signal in RNA-seq due to incomplete RNAP processivity and pervasive transcription regulated by the changing levels of Rho protein observed in different conditions (Bidnenko & Bidnenko, 2018; Wade & Grainger, 2014). These very long predictions can mask predicted transcripts in the same region from other samples, obscuring potentially interesting shorter transcripts expressed in different conditions. For this reason, transcripts longer than 1000 nucleotides were eliminated before combining the predictions between datasets. The predicted annotations for each dataset were combined into a single annotation file, adding the union of the predicted boundaries to the reference genome for H37Rv (AL123456.3). Predictions that overlapped with annotated ncRNAs and UTR predictions that overlapped sRNA predictions from a different dataset were eliminated. Transcript quantification was repeated on each dataset using the resulting combined annotation file and the count data from each dataset was merged into a single counts matrix.

*DESeq2* v1.30.1 (Love et al., 2014) was used on the complete counts matrix including the filtered *baerhunter* predictions to calculate size factors, estimate dispersion and normalise the data with the regularised log transformation function (Supp figures, S1 and S2). The normalised data was checked for potential batch effects using PCA plots and hierarchical dendrograms. *Limma* v3.46.0 (Ritchie et al., 2015) ‘*removeBatchEffect*’ was applied with a single batch argument to remove batch effects associated with the first component (batching the data according to dataset due to technical differences) while preserving differences between samples. The final hierarchical dendrogram, post-batch correction, indicates successful application as samples cluster by similar experimental conditions, rather than by dataset alone (Figure 2 compared to Supp figure S3). Samples from experiment PRJEB65014 continue to group together, but as they represent single replicates in unique conditions, it is difficult to estimate the influence of confounding batch effects for these samples.

**Figure 2.**
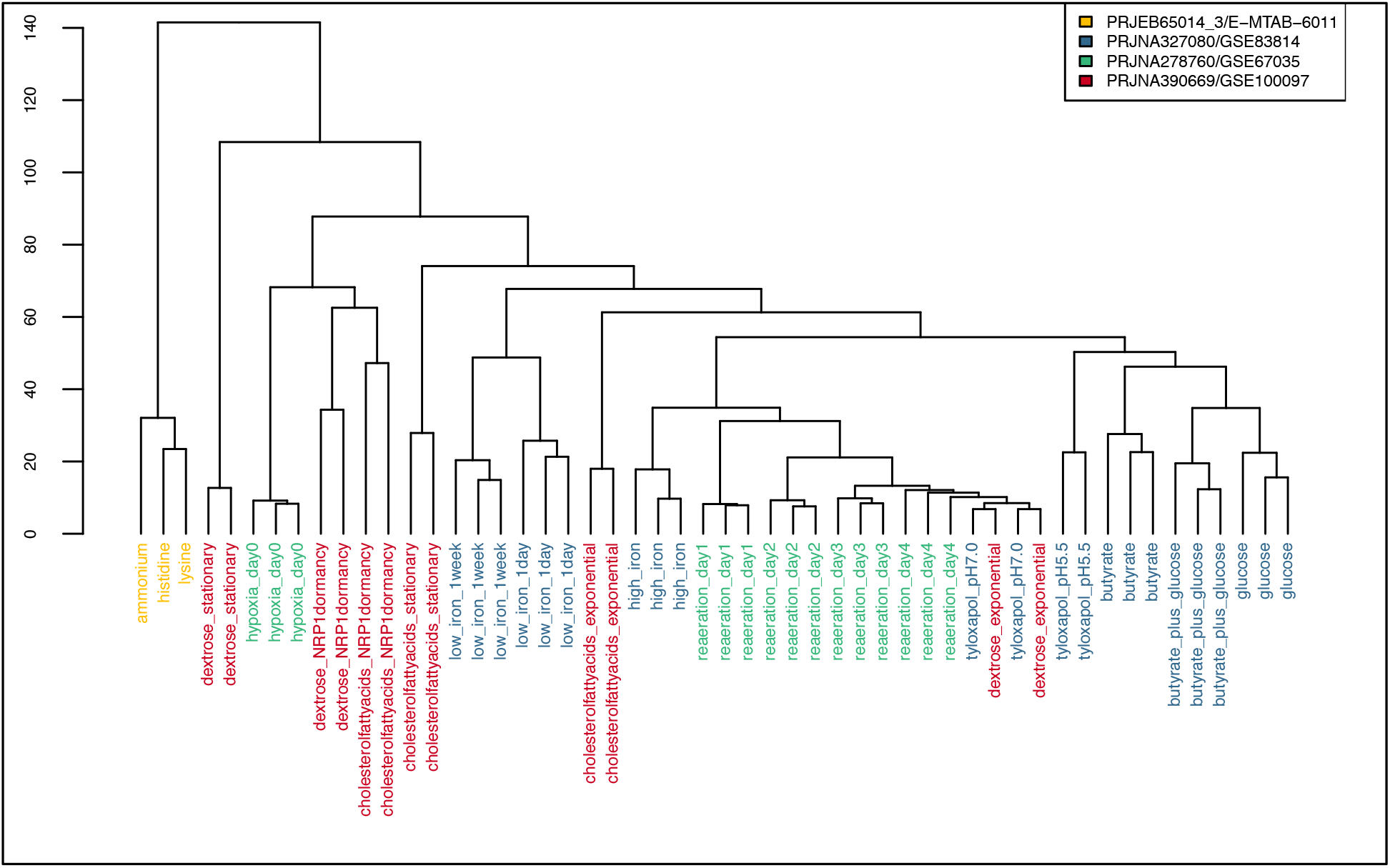
Hierarchical dendrogram of *rlog* transformed and *limma* batch corrected expression data by sample. The sample labels are coloured by dataset, demonstrating that they are clustering by condition, rather than experiment.

### Creation of the WGCNA network

The normalised and batch-corrected expression matrix was used to create a signed coexpression network using the R package, *WGCNA* v1.69 (Langfelder & Horvath, 2008), with the following parameters: corType = “pearson”, networkType = “signed”, power = 12, TOMType = “signed”, minModuleSize = 25, reassignThreshold = 0, mergeCutHeight = 0.15, deepSplit = 2. In this type of network, the ‘nodes’ are the genes, and the ‘edges’, or links, are created when gene expression patterns correlate. In contrast to unweighted binary networks where links are assigned 0 or 1 to indicate whether or not the genes are linked, in a weighted network the links are given a numeric weight based on how closely correlated the expression is. *WGCNA* first calculates the signed co-expression similarity for each gene pair. The absolute value of this correlation is raised to a power (determined by the user, based on a scale-free topology model that mimics biological systems (Supp figure S4) in order to weight the strong connections more highly than the weaker connections. The resulting similarity matrix is used to cluster groups of genes with strong connections to each other in a non-supervised manner (i.e., it doesn’t use any previous information about gene groups or connected regulons). A cluster dendrogram is created (Supp figure, S8) and closely connected branches of the dendrogram are merged into modules based on a cut-off value (also a parameter controlled by the user). The modules are defined by a ‘module eigengene’ (ME), which explains most of the variance in the expression values in the module. The connectivity of the MEs define the shape of the overall network (Supp figure, S9). The modules can then be tested for potential correlations with experimental conditions while reducing the degree of penalties for multiple testing. In signed networks, correlation of the module with a condition can be in either the positive or negative direction, as modules include transcripts that are similar in both the degree and direction of correlation, allowing for a more fine-grained analysis than with unsigned networks (Supp figure, S10).

To test correlations of modules with experimental conditions, the individual RNA-seq samples were assigned to a condition based on the experimental description in the project metadata. Some of these conditions were shared among the different projects, so when appropriate, samples from different datasets were assigned the same condition, resulting in 15 tested conditions. For example, late-stage reaeration samples were tested along with exponential growth samples, and samples that tested hypoxia and cholesterol utilisation together were included in multiple conditions. Models of hypoxia differed between the RNA-seq projects, and these samples were assigned to different conditions: ‘hypoxia’ versus ‘extended hypoxia’ (Supp Table 1, ‘Condition summary’ tab). All correlations were made using robust biweight midcorrelation tests and all p-values were corrected for multiple testing with the Benjamini-Hochberg (BH) method (Benjamini & Hochberg, 1995). Significance was evaluated as an adjusted p-value (p_adj_) of < 0.05.

### Module Enrichment

Modules were interrogated for enrichment for Gene Onotology (GO) terms (Ashburner et al., 2000; The Gene Ontology Consortium, 2021), Clusters of Orthologous Groups (COG) (Galperin et al., 2021), KEGG pathway genes (Kanehisa et al., 2022), functional categories and literature searches for known regulons. GO terms, COG term and KEGG pathway enrichment were accessed programmatically using the DAVID web service (Huang et al., 2009b, 2009a; Jiao et al., 2012) to query the list of protein-coding genes from each module for enrichment. Enrichment was determined using a modified one-sided Fisher’s Exact Test (‘EASE’ score) with BH correction for multiple testing, with p_adj_ < 0.01 considered significantly enriched for a particular term, pathway or COG term. Enrichment for the 11 functional categories from Mycobrowser annotation (Kapopoulou et al., 2011) was determined using a one-sided Fisher’s Exact Test with BH correction for multiple testing. Modules were enriched for a particular functional category if p_adj_ < 0.01. Lists of genes associated with known regulons were mined from literature and enrichment was tested using the same one-sided Fisher’s Exact Test as above with a p_adj_ < 0.01 cut-off for enrichment.

Non-coding RNA prediction, network analysis and subsequent data manipulation was performed with R (v4.0.5, 2021-03-31). All plots were made in R with the following packages: *WGCNA* (v1.69), *dendextend*(v1.15.2), *ggplot2*(v3.3.5). Scripts and expression data are available at **https://doi.org/10.5281/zenodo.7319853.**

## RESULTS AND DISCUSSION

### Mtb expresses an extensive range of ncRNA transcripts over a wide variety of experimental conditions

*Mycobacterium tuberculosis* RNA-seq datasets were selected from publicly available data to find experiments using the wild-type H37Rv strain and representing a range of growth conditions the pathogen may encounter in a host environment. Four datasets passing our quality standards were subjected to our analysis pipeline (see Material and Methods) and included 52 samples under 15 different experimental conditions (Supp Table 1, ‘Samples’ tab). The R package, *baerhunter* (Ozuna et al., 2019), was used to predict ncRNA in intergenic regions, antisense RNA (opposite a protein-coding gene) and UTRs at both the 5’ and 3’ ends of genes by searching the mapped RNA-seq data for expression peaks outside of the annotated regions in the reference sequence for H37Rv. Non-coding RNA predictions from each dataset were filtered for low expression and combined to create a single set of non-overlapping annotations that encompassed all predictions made from any sample under any experimental condition. In total, 1283 putative sRNAs were predicted (including both truly intergenic transcripts as well as those antisense to a protein-coding gene, or annotated RNA) and 1715 UTRs which includes all transcribed regions outside of annotated protein-coding sequences at both 5’ and 3’ ends, as well as the non-coding regions between adjacent genes in operons. All putative ncRNA transcripts (sRNAs and UTRs) were searched for a TSS near the start of the predicted 5’ boundary using previously published annotations (Cortes et al., 2013; Shell et al., 2015). Annotated TSSs were found within 20 nucleotides of the 5’ end in 43% of the predicted sRNA transcripts. Predicted 5’ UTRs had a TSS within 10 nucleotides of the start in 42% of cases, compared with 3% of the predicted 3’ UTRs. Where the UTR covered the entire sequence between two protein-coding regions (labelled as ‘between’ UTRs), 9% had a TSS in the first 10 nucleotides of the sequence (Table 2 and Supp Table 2 ‘putative_sRNAs’, ‘putative_UTRs’ tabs).

**Table 2.**
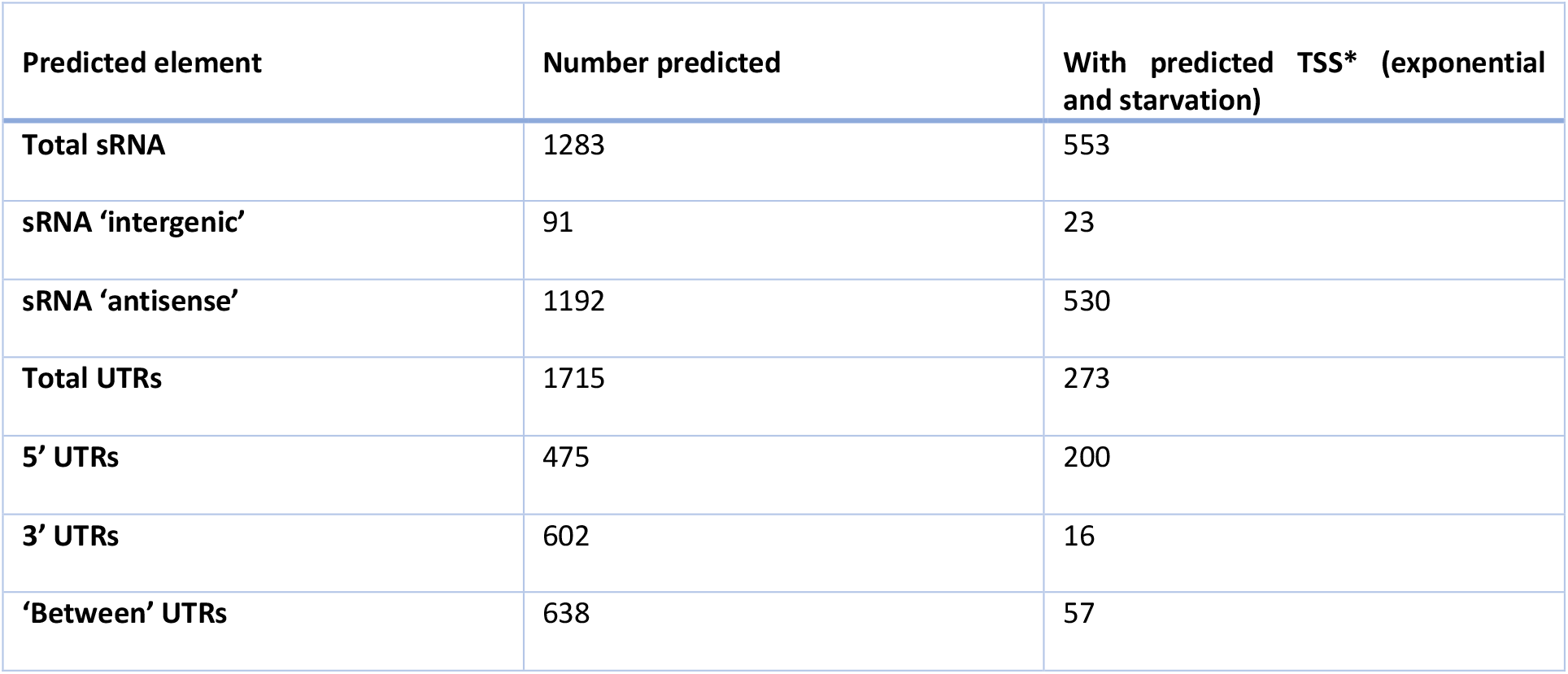
Tally of predicted expressed elements in the *baerhunter-generated* combined annotation file. 4015 protein-coding genes were included in the annotation. ‘Between’ UTRs cover the entire sequence between two protein-coding regions. *TSS predictions from (Cortes et al., 2013; Shell et al., 2015).

| Predicted element | Number predicted | With predicted TSS* (exponential and starvation) |
| --- | --- | --- |
| **Total sRNA** | 1283 | 553 |
| sRNA ‘intergenic’ | 91 | 23 |
| sRNA ‘antisense’ | 1192 | 530 |
| **Total UTRs** | 1715 | 273 |
| **5’ UTRs** | 475 | 200 |
| **3’ UTRs** | 602 | 16 |
| **‘Between’ UTRs** | 638 | 57 |

The predicted sRNAs were further annotated using the accepted nomenclature (Lamichhane et al., 2013) which identifies the putative ncRNA relative to annotated gene loci and differently signifies truly intergenic sRNAs and those that overlap any part of a protein-coding region on the opposite strand. Most of the putative sRNAs are antisense to the protein-coding region of one or more genes, but 91 putative sRNAs have predicted boundaries that do not overlap an annotated transcript on either strand (or overlap an annotated transcript on the opposite strand by fewer than 10 nucleotides). This number is most probably an underestimate of the truly ‘intergenic’ sRNAs in the genome, as many of the sRNA predictions appear over-estimated at the 3’ end, effectively classifying them as an antisense RNA even though the 5’ half of the transcript does not overlap any genes on the opposite strand. Isoforms of annotated sRNAs can be subject to post-transcriptional processing to create an active transcript (Moores et al., 2017) and post-transcriptional processing of 3’ ends *in vivo* is more likely the norm for most prokaryotic transcripts (Wang et al., 2019). However, for our purposes, any RNA-seq transcripts that extend to overlap a protein-coding gene on the other strand in any dataset will be labelled as antisense RNA.

The generated combined annotation file was used to quantify the expression of all 7043 expressed elements, including every annotated CDS, annotated ncRNA and predicted ncRNA, in each sample. Raw counts of expression varied greatly among the datasets due to different sequencing depth, as well as between some samples within datasets (as would be expected with different environmental conditions), and only three protein-coding genes showed no expression in any sample. The raw expression counts were transformed using DESeq2’s rlog function (Love et al., 2014), and plots of the dispersion of count data show that the median expression level between samples and between datasets has been normalised (Supp figures S1, S2). The distribution of the normalised expression levels of protein-coding regions alone shows consistent median expression levels across the entire dataset, however distribution of the normalised data restricted to putative sRNAs shows more variability, with certain conditions showing increased or decreased expression of these transcripts (Supp figures S5-S7). This is not unexpected, given that several studies have identified pervasive transcription in hypoxic infection models, stationary phase and dormancy. This is accompanied by a concomitant increase in non-rRNA abundance (especially antisense RNA transcripts) and in the number of predicted TSSs in Mtb and *M. smegmatis* (a fast-growing, non-pathogenic strain) (Arnvig et al., 2011; Ignatov et al., 2015; Martini et al., 2019).

### Module networks represent groups of co-expressed genes and predicted non-coding RNA

#### Creation of the WGCNA network

A weighted co-expression network was created from the normalised RNA-seq expression data using *WGCNA* (Langfelder & Horvath, 2008) (see Materials and Methods). This program segregates genetic elements with similar patterns of expression over a range of samples into modules. The modules represent sub-networks of connected genes, and functional relationships can be explored among the members of the individual modules. The ‘hub’ genes represent the most highly connected genetic elements within a module and have highest module membership values. Module membership is measured by correlation of the expression of the individual genes with the module eigengene (ME), the vector that best represents the variation in the module.

The signed co-expression network presented in this paper consists of 54 different modules, assigning 99.3% of the expressed elements (CDS, putative UTRs and putative sRNAs) into 53 modules, with 46 unassigned elements clustered in the ‘grey’ module (Supp Table 2, ‘Module_Overview’ tab). Module size ranged from 766 to 25 expressed elements. The modules (using the ME) were tested for correlations with the various conditions used in the RNA-seq experiments (see Materials and Methods). The RNA-seq data was categorised into 15 different experimental conditions in total with varying numbers of replicates (Supp Table S1, ‘Condition Summary’ tab), therefore, a statistically significant correlation of modules with every condition was not expected. However, some modules do show significant correlations with conditions such as iron restriction, cholesterol media, hypoxia and growth phase and this can be informative when considering the association of the gene groups with biological processes (Figure 3).

**Figure 3.**
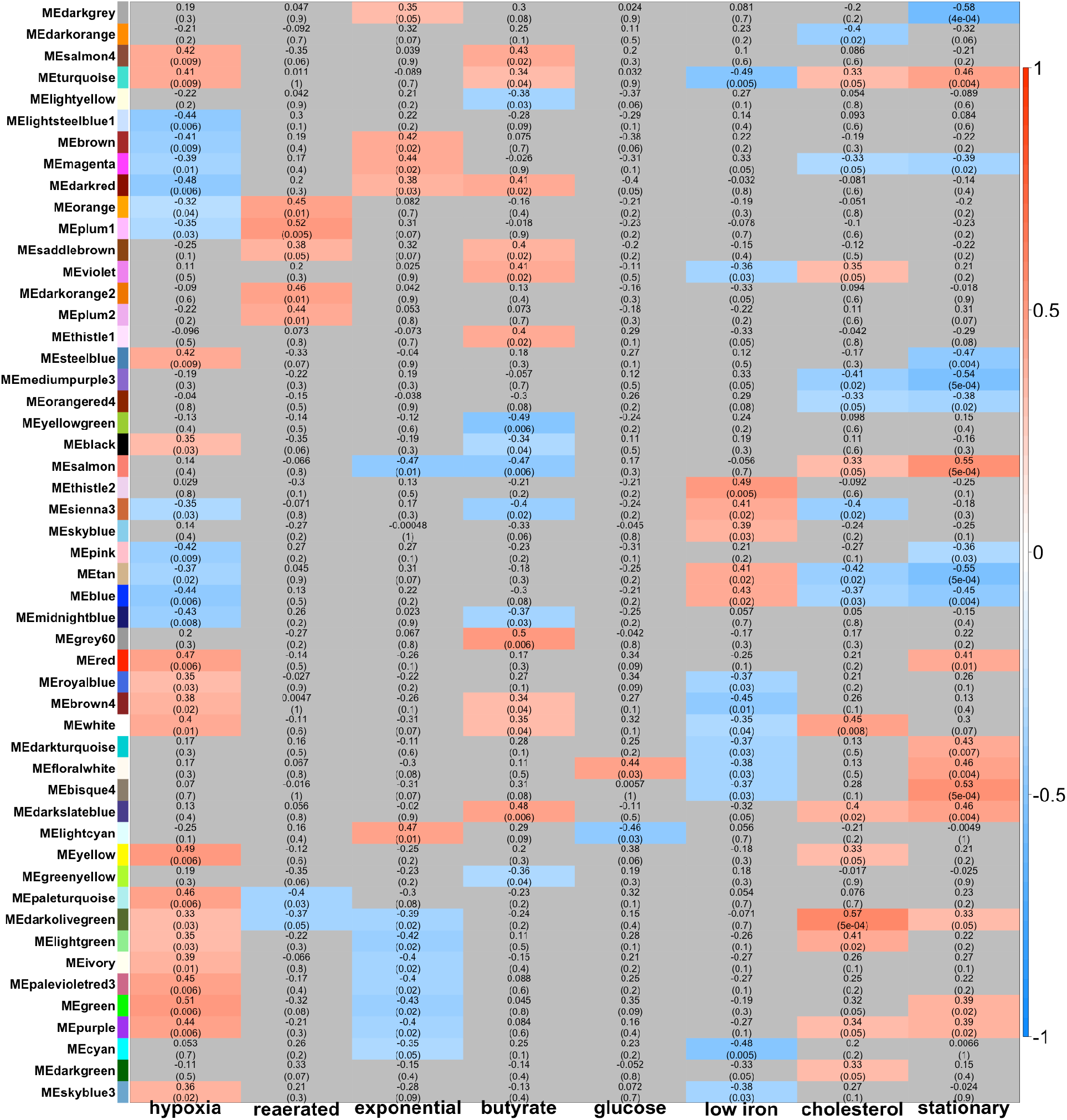
Heat map of correlation of module eigengene (ME) of each module with selected experimental conditions. Correlation was calculated using biweight midcorrelation (bicor) and p-values were adjusted for multiple testing (BH-fdr). Positive correlation is red, negative correlation is blue. Non-significant correlations in grey (p_adj_ > 0.05).

#### Well-established regulons cluster together in single modules

In many cases, the gene membership of the modules includes well-established regulons or groups of functionally related genes, establishing the biological relevance of the module sub-networks and proof of concept for the application of WGCNA on such a heterogenous dataset. For example, the DosR regulon is a well-studied regulon associated with hypoxia and stress responses (Du et al., 2016; Rustad et al., 2008; Voskuil et al., 2004). 47 of 48 previously identified DosR-regulated genes are found in a single module, *‘cyan*’, representing statistically significant enrichment of DosR-regulated genes in the module (one-sided Fisher’s exact test, p_adj_=3.81e-53). The *‘cyan’* module also includes 5 genes from the PhoP regulon which is associated with hypoxic response and coordination with the DosR regulon (Gonzalo-Asensio et al., 2008; Singh et al., 2020) and the DosR-regulated ncRNA, DrrS/MTS1338, known to be upregulated in hypoxic conditions (Ignatov et al., 2015; Moores et al., 2017). Unsurprisingly, the *‘cyan’* ‘module is enriched for the GO term, ‘response to hypoxia’, however, a statistically significant correlation was not seen with the hypoxia condition (though it is negatively correlated with the exponential growth condition, bicor=-0.35, p_adj_=0.05) (Figure 3). The KstR regulon includes 74 genes under control of the TetR-type transcriptional repressor, KstR, known to be involved in lipid catabolism and upregulated during infection (Kendall et al., 2007, 2010; Nesbitt et al., 2010). The *‘royalblue’* module is significantly enriched for known KstR-regulated genes (one-sided Fisher’s exact test, p_adj_ = 5.06e-30) with 30 of 72 KstR-regulated genes clustering together in the module. This module is enriched for genes of the KEGG pathway for steroid degradation (p_adj_= 3.32e-10) and the GO term ‘steroid metabolic process’ (p_adj_ = 5.62e-16). The module shows statistically significant positive correlation for hypoxia (bicor=0.35, p_adj_=0.03) and negative correlation with the low iron condition (bicor=-0.37, p_adj_=0.03) (Figure 3). Genes involved in mycobactin synthesis are nearly all found in the *‘grey60’* module (one-sided Fisher’s Exact test, p_adj_= 1.23e-17), a module enriched for the KEGG pathways ‘siderophore metabolic processes’ and ‘arginine biosynthesis’. As these examples show, known associated genes are co-located in modules which represent a functional group of genes that have co-regulated expression under various experimental conditions. The modules can be further explored to identify novel associations.

#### Predicted non-coding RNAs are enriched in certain modules

Putative sRNAs and/or predicted UTRs were distributed throughout all modules in the network (Figure 4, Supp Table 2, ‘Module_Overview’ tab). The number of predicted sRNAs were statistically enriched in seven modules and predicted UTRs enriched in another seven modules (one-sided Fisher’s exact test, p_adj_ < 0.01, Supp Table 2, ‘Module_Overview’ tab). A roughly linear relationship between the number of CDS and the number of UTRs, is to be expected, given that UTRs are defined by the *baerhunter* algorithm by their position at the start or end of protein-coding genes (Ozuna et al., 2019). However, if the UTRs are positioned in an operon, there will be a smaller increase in the relative number of UTRs with an increasing number of protein-coding genes, as UTRs between two protein-coding genes are predicted as a single UTR. As expected, the two modules that include the highest number of predicted operons (from OperonDB, Chetal & Janga, 2015), *’turquoise’* and *‘brown’*, have a lower relative proportion of UTRs; however, the *‘blue’* module, which includes 15 complete predicted operons, is significantly enriched for UTRs (p_adj_ = 6.79e-21) (Figure 5).

**Figure 4.**
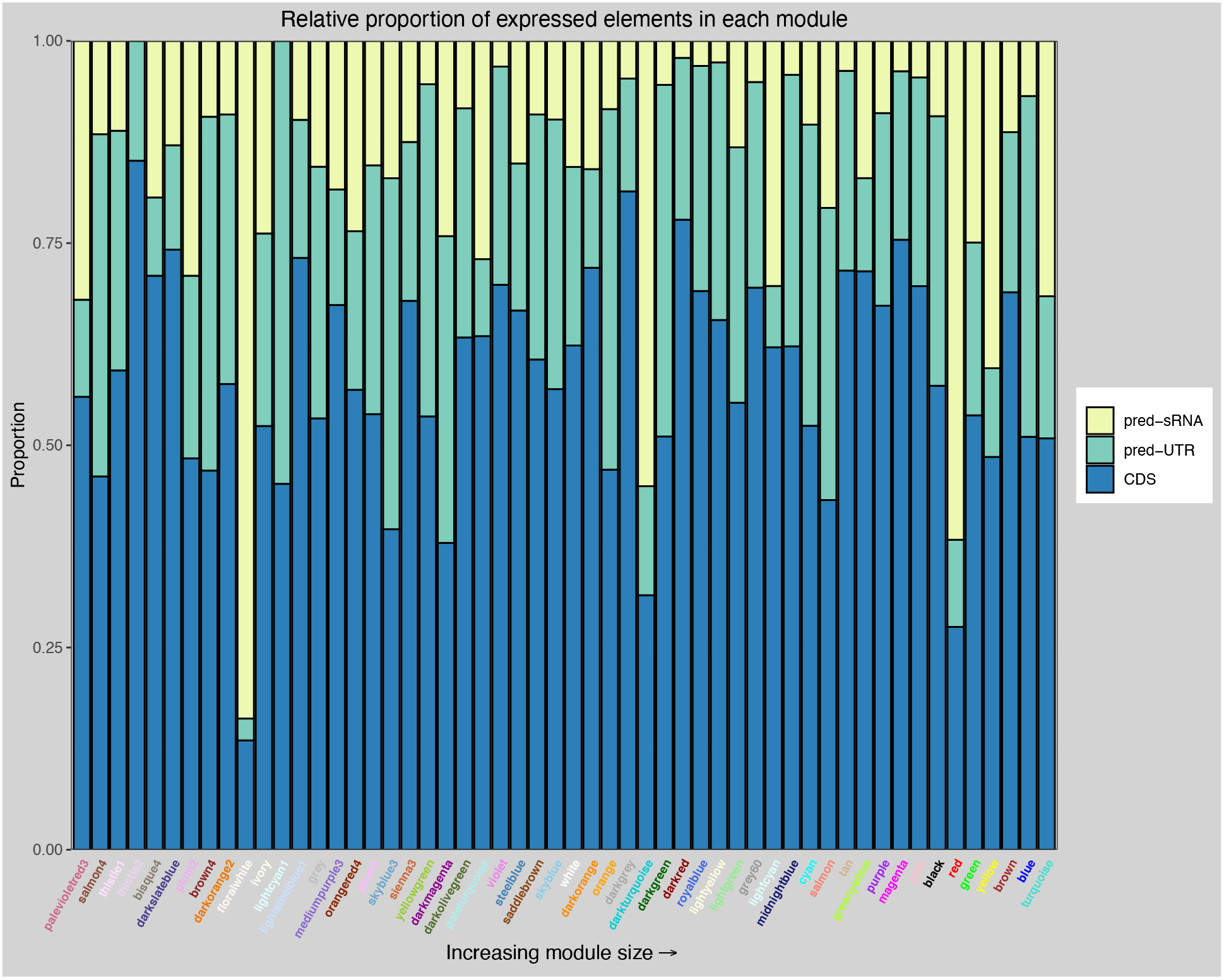
Relative proportion of annotated CDS, predicted UTRs and predicted sRNAs in each module, ordered by module size.

**Figure 5.**
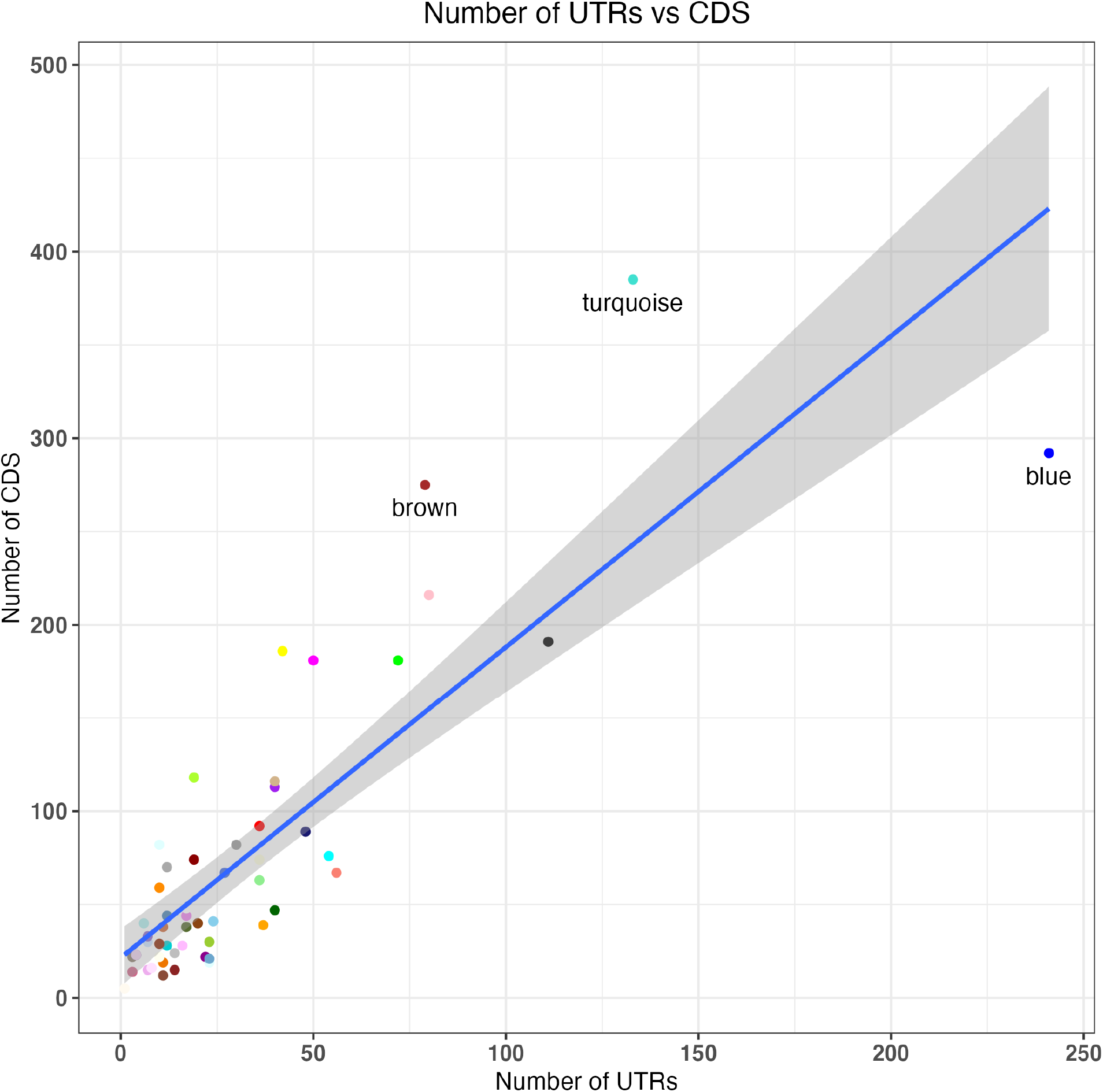
Plot of number of UTRs against number of CDS in each module. Grey shading indicates confidence interval of 0.95.

Within the module sub-networks, the tight co-expression of protein-coding genes and ncRNA is reflected by the number of ncRNA found among the most connected elements in the module. The ‘hub’ elements are those with the best correlation to the ME and therefore the most tightly connected elements in the individual module networks. In 14 modules, ncRNA (both predicted and annotated) make up more than half of the elements with module membership values (MM) > = 0.80 (our threshold for identifying hub elements) (Supp Table 2, ‘Hub_info’ tab). These associations may implicate ncRNA as co-conspirators in regulatory pathways implemented to adapt to conditions such as hypoxia, cholesterol media and low iron. The 30 annotated ncRNAs in the Mtb reference genome (AL123456.3) are spread over 20 modules, with 10 of them hubs of the module, and one unassigned (‘grey’ module) (Supp Table 2, ‘Annotated ncRNA’ tab). For example, Ms1/MTS2823, observed to be the most abundantly expressed ncRNA in expression studies over various stress conditions (Arnvig et al., 2011; Arnvig & Young, 2012; Ignatov et al., 2015; Šiková et al., 2019), is a hub element in a module that is positively correlated with cholesterol-containing media conditions (*‘darkgreen’*, bicor=0.35, p_adj_=0.04) (Figure 3). This module is significantly enriched for KEGG pathways, including: Pyruvate metabolism (p_adj_ = 3.1e-3) and two-component systems (p_adj_ = 3.8e-3), and GO terms: plasma membrane respiratory chain complex II and plasma membrane fumarate reductase complex. Mcr7/ncRv2395A, found to be part of the PhoP regulon (Solans et al., 2014), is a hub in the *‘violet’* module enriched for lipid metabolism and PE/PPE functional categories, correlated positively with growth in cholesterol (bicor= 0.35, p_adj_= 0.04) and butyrate (bicor= 0.41, p_adj_= 0.02) and negatively correlated with low iron (bicor= −0.36, p_adj_= 0.03) (Figure 3). F6/ncRv10243/SfdS, a sRNA upregulated in starvation and mouse infection models, is thought to be involved in regulating lipid metabolism and long-term persistence (Houghton et al., 2021). This ncRNA is a hub in a module found to be enriched in ‘lipid metabolism’ genes (*‘saddlebrown’*) and found to be correlated positively with reaerated culture (bicor= 0.38, p_adj_= 0.04) and butyrate (bicor= 0.4, p_adj_= 0.02) conditions (Figure 3).

#### UTR and adjacent ORF expression differ in over 50% of cases

We were interested to see how many of the predicted UTRs were assigned the same module as the adjacent ORF—indicating whether the ORF and its adjacent UTR were co-regulated. Intuitively, the UTR of a protein-coding gene would be expected to be expressed as a single transcript along with the ORF and show similar expression patterns. However, both 5’ and 3’ UTRs can act independently of the attached ORF and RNA abundance in RNA-seq experiments reflects both transcription activity and transcript stability. For example, some 5’ UTRs are known to contain regulatory elements, such as riboswitches, that alter the transcription of the downstream ORF (Dar et al., 2016; Kipkorir et al., 2021; Schwenk & Arnvig, 2018; Warner et al., 2007), whereas sRNAs cleaved from 3’ UTRs have been shown to regulate the stability of the remaining transcript--with different half-lives as a result (Chao et al., 2012; Dar & Sorek, 2018; Menendez-Gil & Toledo-Arana, 2021). Of the *baerhunter* -predicted UTRs labelled 5’ and 3’, the UTRs co-segregated with the ORF they were closest to in fewer than half of cases (Table 3). We would expect correctly-identified 5’ UTRs to utilise a TSS (whether or not there is a known predicted TSS), whereas it appears functional 3’ UTRs are more likely to be cleaved from the longer mRNA transcript (Dar & Sorek, 2018; Menendez-Gil & Toledo-Arana, 2021; Ponath et al., 2022). Our data confirms this: transcripts classified as 5’ UTRs are much more likely to have a predicted TSS in the first 10 nucleotides than transcripts classified as 3’ UTRs (42% vs 2.7%). Approximately 9% of the UTRs predicted to be between ORFs (labelled, ‘Between’ UTRs) have predicted TSS (Table 3). The presence of a TSS in the first 10 nucleotides of the predicted UTR appeared to have little bearing on whether or not the UTR and its adjacent ORF are assigned to the same module, with 43% of 5’ and 19% of 3’ UTRs with a predicted TSS co-assigned with their adjacent ORF partner. 42% of the ‘Between’ UTRs do not segregate with either the ORF upstream or downstream, indicating their expression is, to some degree, independent of either adjacent ORF. 195 UTRs were found to be hubs in modules independent of their adjacent ORF(s), with 27 including a predicted TSS. All ‘independent’ UTRs are found in Supplementary Table 2, ‘independent_UTRs’ tab.

**Table 3.** UTRs and module assignment of adjacent ORFs. DS=downstream, US=upstream. TSS indicates presence of annotated TSS in first 10 nucleotides of predicted UTR (Cortes et al., 2013; Shell et al., 2015).

|  | Total (excluding grey) | Number with TSS | Number in same module as adjacent ORF | Proportion of UTRs in same module as adjacent ORF |
| --- | --- | --- | --- | --- |
| **5' UTR** | 471 | 198 | 173 DS | 37% |
| **3' UTR** | 597 | 16 | 254 US | 43% |
| **BTWN UTR** | 633 | 56 | 112 DS<br>116 US<br>137 both | 18%<br>18%<br>22% |

#### Antisense RNAs are hubs in modules independent of cognate ORF

It has been observed that the overall abundance of antisense RNA and other non-ribosomal RNA increases upon exposure to stress such as hypoxia and nutrient restriction (Arnvig et al., 2011; Ignatov et al., 2015), and in our network, ncRNA are well-connected in various modules that include known transcription factors and gene regulons associated with stress responses. Not unexpectedly, very few (5%) of the predicted antisense transcripts were assigned to the same module as the protein-coding region overlapping on the opposite strand (choosing the most downstream locus in the event of multiple overlapping ORFs), signifying distinct patterns of expression for transcripts on opposite strands, possibly due to independent or bi-directional promoters and/or overlapping transcription termination sites. Bi-directional promoters have been identified in multiple prokaryotic genomes, and competition for RNA polymerase (RNAP) binding among divergently transcribed sense/antisense pairs may function as a mechanism for regulation of gene expression (Ju et al., 2019; Warman et al., 2021). Long 3’ UTRs that overlap with converging protein-coding genes on the opposite strand (or with the 3’ UTR) can create an ‘excludon’ regulatory arrangement, where transcription of the two opposite mRNAs is simultaneously regulated by RNase targeting, or mutually exclusive due to RNAP collision (Sáenz-Lahoya et al., 2019; Toledo-Arana & Lasa, 2020). Examining the module groupings of the antisense RNAs and their base-pairing target on the other strand may provide insight on which genes are regulated by antisense transcription.

### Focus on Selected Module Networks

The large-scale transcription analysis presented here is useful for the more global analysis of the overall trends related to ncRNA and transcription, but there is a great deal of information to be gleaned by more fine-grained inspection of individual module groupings. To discover novel associations in such a large and complex dataset, we have selected a few modules for closer examination, focussing on those that contain gene groups or regulons related to the tested conditions. Many of the modules that contain interesting correlations or gene regulon enrichments also include an abundance of putative sRNAs and UTRs. Using the ‘guilt by association’ principle, we can hypothesise that the well-connected ncRNAs found among the module hub elements have a role in transcriptional ‘remodelling’ in response to changes in environmental conditions such as growth on cholesterol-containing media, restricted iron or hypoxia.

#### Detoxification-linked proteins cluster in the module best correlated with cholesterol media condition

The *‘darkolivegreen’* module showed positive correlation with the cholesterol media condition (bicor=0.57, p_adj_=5.0e-04) and negative correlation with low iron (bicor = −0.48, p_adj_ = 0.001) (Figure 3). Many protein-coding genes involved in detoxification pathways are hubs in the module, including several encoding transmembrane proteins such as the *mmpL5-mmpS5* efflux pump operon (Rv0676c-Rv0677c), as well as the next gene downstream, Rv0678, which was identified as part of a ‘core lipid response’ in differential expression analysis in lipid-rich media (Aguilar-Ayala et al., 2017). The 5’ UTR for Rv0677c and 3’ UTRs for Rv0676c and Rv0678 are also hubs. This operon is involved in siderophore transport and expressed in cholesterol and lipid-rich environments (Aguilar-Ayala, et al., 2017; Pawełczyk et al., 2021). The module contains several Type II toxin-antitoxin systems including VapBC12 (Rv1720c1721c), VapBC41 (Rv2601A-2602), RBE2 (relFG, Rv2865-2866) and vapB36 and vapB40 which may have roles in adaptation to cholesterol and the evolution of persisters (Ramage et al., 2009; Sala et al., 2014). VapBC12, specifically, has been shown to inhibit translation and promote persister phenotypes in response to cholesterol (Talwar et al., 2020). Other detoxification-linked genes in the module, such as the ABC-family transporter efflux system, Rv1216c-1219c, have also been implicated in transcriptomic remodelling in response to cholesterol (Aguilar-Ayala et al., 2017; Pawełczyk et al., 2021).

Two adjacent predictions, the 3’ UTR for Rv1772 (putative_UTR:p2006948_2007063) followed by ncRv1773/putative_sRNA:p2007213_2007377, are hubs in the *‘darkolivegreen’* module. Together, they extend to overlap the antisense strand of a large portion of Rv1773c, a probable transcriptional regulator in the IclR-family, found in a different module (*‘turquoise’*). The 3’ UTR for Rv1772 has been previously identified as an abundant antisense transcript during exponential growth (Arnvig et al., 2011). The start of the predicted sRNA transcript has no known TSS and could instead be an extension of the predicted 3’ UTR (Supp figure S11). (When combining predicted annotations from different datasets, long predicted UTRs that overlapped shorter sRNA predictions were discarded, see Methods). In *E.coli*, the IclR-family transcriptional regulators demonstrate both activating and repressing activities on targets such as multidrug efflux pumps and the *aceBAK* operon which regulates the glyoxylate shunt (Zhou et al., 2012). *Icl2a* (Rv1915) is one of the Mtb isoforms of the isocitrate/methylocitrate lyase gene, *aceA*, and may be regulated by Rv1773c, as seen in *E.coli*. Icl2a, Rv1772, its predicted UTR and the antisense RNA (ncRv1773) are all hubs in the *‘darkolivegreen’* module. *Icl2a* has been observed to be upregulated with cholesterol as the sole carbon source and likely has a second function as part of the methylcitrate cycle to convert the fatty acid metabolites propionate and propionyl CoA to less toxic compounds (Bhusal et al., 2017; Pawełczyk et al., 2021).

#### Module correlated with reaeration after non-replicating persistence includes genes for amino-acid synthesis and cell wall remodelling

The module, *‘saddlebrown’* is enriched for GO-terms for various amino-acid metabolic processes and COG ‘lipid metabolism’. It is positively correlated with reaeration after non-replicating persistence (bicor= 0.38, p_adj_= 0.04) and butyrate-containing media (bicor= 0.4, p_adj_= 0.02) (Figure 3). This pairing of upregulation of amino-acid synthesis and upregulation of the synthesis of cell wall lipids has been observed in the ‘lag phase’ after reaeration for increased protein synthesis (Du et al., 2016). The hubs of the *‘saddlebrown’* module include several predicted sRNAs, and the annotated sRNA, F6. F6/ncRv10243/SfdS is a sigF-dependent ncRNA which has been shown to be induced in nutrient starvation, oxidative stress, acid stress (Arnvig & Young, 2009; Houghton et al., 2021) and the fatty acid hypoxia model (Del Portillo et al., 2019). In addition to being expressed from its own promoter, F6/SfdS has been proposed to be co-transcribed with the upstream gene *fadA2* (Rv0243), a probable acetyl-CoA acyltransferase; however, *fadA2* is clustered in a different module from SfdS (‘*darkred*’).

One of the predicted sRNAs among the *‘saddlebrown’* module hubs is antisense transcript ncRv2489/putative_srna:p2801108_2801678 with a TSS at 2801108. This overlaps the 3’ end of PE-PGRS43 (Rv2490c) (Figure 6). There is a short reading frame (30 nucleotides, 10 amino acids) initiating from a Methionine at this TSS that suggests a possible dual-function sRNA or sORF with independent function. A shorter, possibly-leadered, sORF was predicted by Shell et al. (2015) that falls within this region (2801238..2801261). The TSS for the predicted sRNA overlaps the 5’ end of Rv2489c, a short, hypothetical ‘alanine-rich protein’. The TSSs for these convergently overlapping transcripts are 42 nts apart and may involve RNAP collision if both are transcribed simultaneously. Therefore, transcription of the predicted sRNA could impact either Rv2489c and/or PE-PGRS43 expression through two different mechanisms. Another hub sRNA in *saddlebrown’* includes ncRv1450/putative_sRNA:p1630466_1631246, which has a TSS at 1630466 and is likely to be an intergenic transcript between two divergently transcribed genes on the opposite strand, tkt (Rv1449c) and PE-PGRS27 (Rv1450c), both of which are assigned to different modules. The 3’ end of the prediction includes possible run-on transcription antisense to the 3’ end of PE-PGRS27.

**Figure 6.**
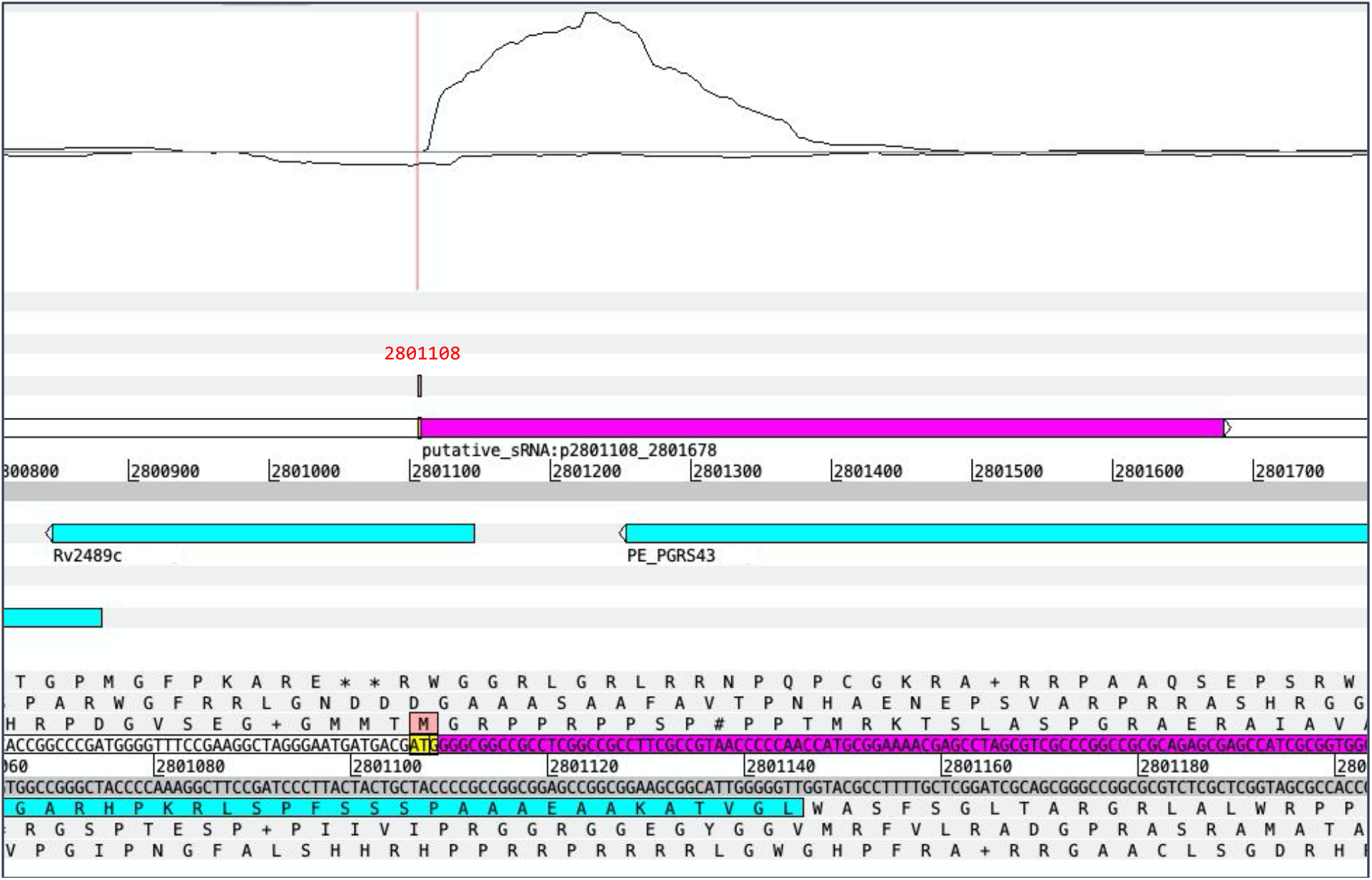
Antisense sRNA, ncRv2489/putative_srna:p2801108_2801678, (magenta bar) overlaps two transcripts and may encode a short peptide. TSS for sRNA indicated in red and corresponding amino acid highlighted in pink. Sample SRR5689230 from PRJNA390669, exponential growth on cholesterol and fatty acid media. Strand coverage using the ‘second’ read of each pair mapping to the transcript strand, visualised using Artemis genome browser (Carver et al., 2012).

The fatty-acid desaturase gene, Rv3229c (*desA3*) is a hub in the module, but without its operon partner, Rv3230c. However, the module does contain an antisense sRNA in this region, ncRv3230/putative_sRNA:p3607084_3607499 which is antisense to the 3’ end of Rv3230c, but lacks a known TSS. Interestingly, Rv3230c has an internal transcription termination site predicted at 3607550 which coincides with the 3’ end of the antisense sRNA (D’Halluin et al., 2022) (Supp figure S12). Another hub antisense sRNA, putative_sRNA:p3608313_3608866/ncRv3231c, overlaps the 3’ end of the upstream gene, Rv3231c, and has a predicted TSS at 3608313.

#### Slow-growth correlated module is associated with transcriptional remodelling and metal ion homeostasis and enriched for sRNAs

The *green’* module contains genes that are associated with transcriptional remodelling in response to hypoxic or stationary growth conditions. It is positively correlated with hypoxic (bicor=0.49, p_adj_=0.004) and stationary (bicor=0.4, p_adj_=0.01) growth conditions, negatively correlated with exponential growth (bicor=-0.44, p_adj_=0.01) (Figure 3) and is enriched for GO terms related to response to metal ions as well as regulation of gene expression. The *‘green’* module contains at least 30 known transcription factors, with 14 of them hubs in the module, including FurA, Zur and sigma factor, SigH, as well as being enriched for SigH regulon genes. Three of the most well-connected transcription factors (furA, smtB and zur) are involved in iron uptake and utilisation, and the Zur-regulated ESAT-6 secretory proteins, esxR and esxS (Rv3019c, Rv3020c), are also present in the module, linking metal homeostasis with response to hypoxia (Maciąg et al., 2007; Zhang et al., 2020). Two chaperonin protein targets of the non-coding RNA F6/Sfds, GroES (Rv3418c) and GroEL2 (Rv0440) are in the module, as well as the chaperonin protein, hsp (Rv0251c), all of which are part of the phoPR virulence-regulating system (Gonzalo-Asensio et al., 2008, 2014).

The *‘green’* module is enriched for sRNAs (p_adj_=0.011). Among the best-connected, are 27 predicted antisense RNAs. One of these hubs, putative_sRNA:p1404640_1404929/ ncRv1257 is antisense to the 3’ end of Rv1257c, a probably oxidoreductase, and another (putative_sRNA:p1771044_1771498/ ncRv1546) is antisense to the 5’ end of a trehalose synthetase, *treX*. Both of these sRNAs have TSSs and are expressed differentially among the tested conditions. Control of reactive oxygen species and synthesis of trehalose intermediates are important for cells in to survive in hypoxic conditions (Eoh et al., 2017; Harold et al., 2019) and antisense RNA may be involved in fine-tuning these responses.

Another antisense RNA, ncRv1358c (putative_sRNA:m1530046_1530745) has a TSS near its start and is found antisense to Rv1359. Rv1359 and the upstream gene, Rv1358, on the opposite strand are very similar to each other (43.7% identity in 197 aa overlap) and to another gene elsewhere in the genome, Rv0891c (48.5% identity in 204 aa overlap) (Kapopoulou et al., 2011). All three genes are possible LuxR family transcriptional regulators which are thought to be involved in quorum-sensing adaptations and contain a probable ATP/GTP binding site motif (Chen & Xie, 2011; Modlin et al., 2021) and are found in different modules. Expression of this antisense sRNA appears to suppress the expression of the transcript on the opposite strand to varying degrees in all conditions (Figure 7). In the cholesterol and fatty acid media samples, expression of a shorter transcript appears to begin inside the Rv1359 ORF, where the transcript is not overlapped by the antisense transcript, possibly utilising an internal TSS at 1530774.

**Figure 7.**
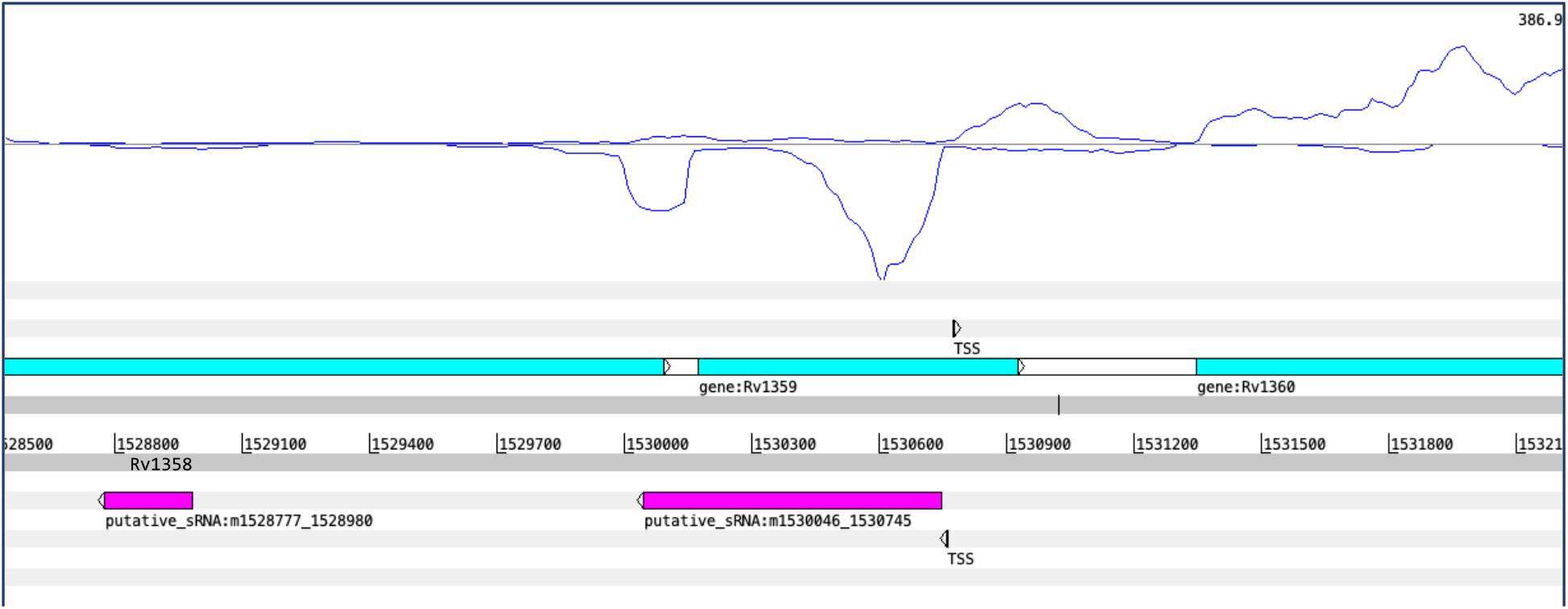
Expression of antisense transcript putative_sRNA:m1530046_1530745 (magenta bar) seems to suppress the expression of most of Rv1359 and Rv1358 in cholesterol and fatty acid media. An internal TSS exists inside the Rv1359 CDS at 1530774 near where expression begins. Note, prediction of an individual sRNA is an aggregate of predictions under different conditions, so will not always match the expression of the sRNA in any particular sample. Sample SRR5689230 from PRJNA27860. Strand coverage using the ‘second’ read of each pair mapping to the transcript strand, visualised using Artemis genome browser (Carver et al., 2012).

### Metal ion homeostasis genes cluster in module that is negatively correlated with the hypoxia condition

The *‘darkrec’* module is negatively correlated with the hypoxia condition (bicor=-0.46, p_adj_=0.005, Figure 3). This module contains most of the ESX-3 genes (Rv0282-Rv0292) related to siderophore-mediated iron (and zinc) uptake in Mtb (Serafini et al., 2013; L. Zhang et al., 2020), with nine of these representing hubs in the module. The module is enriched for the PE/PPE functional category, and includes the two genes preceding the ESX-3 genes, Rv0280 (PPE3) and Rv0281 (a possible S-adenosylmethionine-dependent methyltransferase involved in lipid metabolism, though its position in the genome would suggest regulation could be linked to ESX-3 (Lunge et al., 2020)), and an ESX-5 gene, Rv1797 (*eccE5*). The module also contains another Zur-regulated gene, Rv0106, which is a potential zinc-ion transporter (Zondervan et al., 2018). Among the hubs of the module are several genes related to lipid metabolism and fatty acid synthesis, including: probable triglyceride transporter, Rv1410; the operon consisting of Rv0241c (*htdX*), Rv0242c (*fabG4*), and Rv0243 (*fadA2*) (Dutta, 2018); and a gene involved in the pentose phosphate pathway, *zwf2* (Rv1447c).

There are some well-connected ncRNAs in the *‘darkrec’* module, including a predicted antisense RNA to Rv0281, ‘ncRv0281c’ (putative_sRNA:m341328_342075). This putative sRNA has a predicted TSS at the 5’ end and is transcribed divergently from Rv0282 (*eccA3*). This is one of the rarer cases where the antisense transcript and cognate proteincoding gene (Rv0281) are clustered in the same module. The prevailing direction of transcription at this locus may be a result of competition for RNAP binding at a bidirectional promoter in the predicted 5’ UTR of Rv0282 which also clusters in the module. There are several UTRs in the module hubs, including a 3’ UTR for the gene Rv1133c, *metE* (also found in the module). This UTR was previously identified as abundantly expressed in exponential culture (Arnvig et al., 2011). There is a 3’ UTR for Rv0292 (*eccE3*, also a hub in the ‘*darkred*’ module) that is antisense to a large part of the 3’ end of Rv0293c which has a converging orientation to Rv0292 (Supp figure S13). Rv0293c is a hub in a different module (*‘lghtgreen’*) along with its 3’ UTR. Overlapping 3’ ends of genes could function to regulate transcription, possibly by bi-directional termination brought about by RNAP collision, or function post-transcriptionally by influencing transcript stability (Ju et al., 2019; Vargas-Blanco & Shell, 2020).

#### Module enriched for sRNAs and PE/PPE genes is correlated with stationary condition

The *darkturquoise’* module is enriched with sRNAs, with 33 hub sRNAs. It is negatively correlated with the low iron condition (bicor = −0.37, p_adj_=0.03) and positively correlated with stationary growth (bicor= 0.43, p_adj_=0.007). The genes of the module are enriched for the PE/PPE functional category and there are several PE/PPE genes among the hubs. The previously annotated ncRNA, B11 (also known as 6C or ncRv13660c), is one of the most well-connected elements in the module and overexpression of B11 in *M.smegmatis* has been shown to cause growth arrest and downregulation of a large set of genes including those involved in cell division and virulence, including all the ESX-1 secretion system genes (Mai et al., 2019). Mcr11 is also found in the module. This sRNA is known to respond to the second messenger 3’,5’-cyclic adenosine monophosphate and has been found to be expressed in hypoxic Mtb cultures and in a mouse infection model (Girardin & McDonough, 2020). Mcr11 regulates the expression of several genes that adapt central carbon metabolism during slow growth conditions (Girardin & McDonough, 2020).

There are two well-connected intergenic sRNAs predicted in the *‘darkturquoise’* module. Putative_sRNA:p1164036_1164162 / ncRv11040 is located between PE8 and a possible transposase, Rv1041c, but on the antisense strand. There is a predicted TSS at 1163697, 39 nucleotides upstream of the predicted start sequence. This transcript is in a converging orientation to the transposase and may be instrumental in regulating horizontal gene transfer (Ellis & Haniford, 2016; Lejars et al., 2019). The other intergenic hub is also upstream from possible transposase, Rv3114, but in diverging orientation on the opposite strand. The TSS is at 3481459, and the sRNA is within a predicted MT-complex-specific’ genomic island associated with virulence genes (Becq et al., 2007). Rv3112-14 are clustered in the *salmon’* module.

There are several interesting ‘independent’ UTRs that are well-connected in the module, but their assumed transcriptional partner clusters in another module. There are several predicted TSS’s and transcriptional termination sites (TTS) (D’Halluin et al., 2022) within the predicted boundaries of a 3’ UTR for the gene Rv0281c (putative_UTR:m2337218_2338064) and a predicted sORF based on ribosome profiling (Smith et al., 2022) (Figure 8). This 3’ UTR is antisense to Rv2080, *lppJ*, which is in the *‘cyan’* module along with most of the DosR regulated genes. The higher level of expression of the antisense transcript relative to the accepted sense strand has invited speculation that Rv2080 has been misannotated (K. B. Arnvig et al., 2011). The 5’ UTR of Rv0281c is also a hub in the module and contains predicted TSSs, TTS and sORF. It would be interesting to discover whether these UTRs could have dual functions as regulatory RNA elements as well as being translated into short peptides. Rv2081c is a conserved membrane protein containing a simple sequence repeat of 8 C’s and has been identified as a source of sequence variation in Mtb sputum and culture (Shockey et al., 2019; Sreenu et al., 2007).

**Figure 8.**
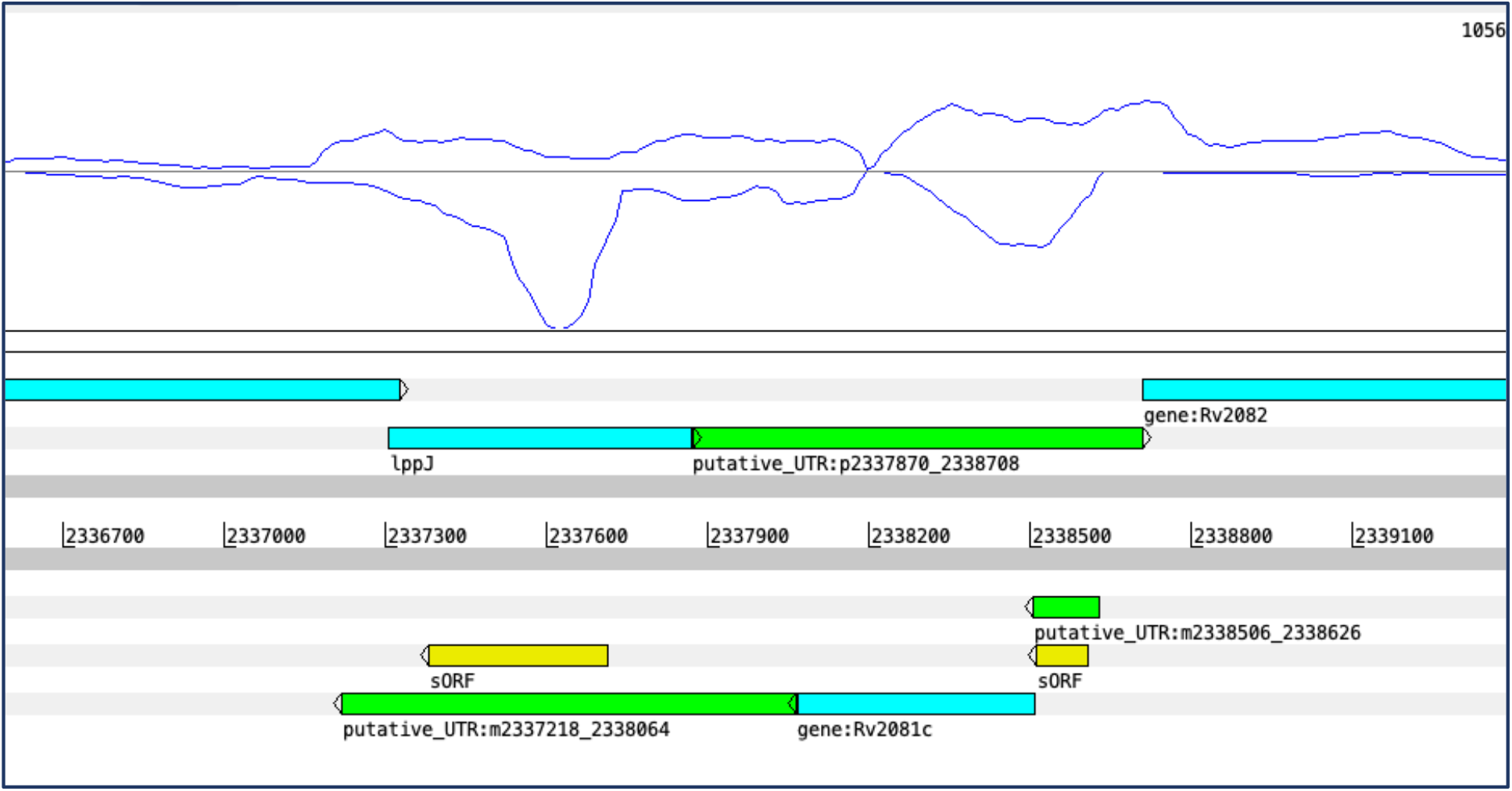
5’ and 3’ UTRs for Rv2081c (green bars) are overlapped by predicted sORFs (yellow bars). (Cortes et al., 2013; Smith et al., 2022). Shown is sample SRR5689224, hypoxia condition from PRJNA27860. Strand coverage using the ‘second’ read of each pair mapping to the transcript strand, visualised using Artemis genome browser (Carver et al., 2012).

The best connected elements in the module are antisense sRNAs, including putative_sRNA:p2081553_2082178/ ncRv1835, with a predicted TSS at its start. It is antisense to Rv1835c, the gene for a putative serine esterase clustered in the *‘mediumpurple3’* module, in particular to the 3’ end of the peptidase domain (Xaa-Pro dipeptidyl-peptidase-like domain) (Blum et al., 2020). Putative_sRNA:m2497549_2498369/ncRv2225c, with a TSS at 2498368, is antisense to Rv2225, coding for a 3-methyl-2-oxobutanoate hydroxymethyltransferase PanB. This gene clusters in the in *‘turquoise’* module.

## CONCLUSION

This paper presents a large-scale network analysis of over 7000 transcripts expressed by Mtb under a variety of conditions. The modules group together clusters of co-expressed protein-coding genes, as well as ncRNA transcripts predicted from RNA-Seq signals. Several modules are statistically enriched for sRNAs, especially those modules positively correlated with hypoxia. The abundance of antisense RNA in conditions of stress has been widely observed, and it is therefore not a surprise to find them in the hubs of these modules. However, it is noticeable that the complementary ORF is usually excluded, which leads us to seriously consider antisense transcription as part of strategic regulation of protein production in response to environmental cues through mechanisms of divergent transcription, translational control or by regulating mRNA stability (Vargas-Blanco & Shell, 2020; Warman et al., 2021). If these strategies actually differ among the members of the MTBC, it may have implications for host specificity and virulence (Dinan, Adam M. et al., 2014). By the same logic, 3’ UTR transcripts clustering in modules distinct from their upstream ORF implies independent function from the ORF. sRNAs generated from 3’ UTRs have been reported in other prokaryotes and evidence points to widespread mRNA processing that could release independent transcripts at the 3’ end (Dar & Sorek, 2018; Desgranges et al., 2021; Updegrove et al., 2019; Wang et al., 2019). In compact bacterial genomes, 3’ UTRs are also found to overlap other 3’ UTRs in a converging transcription pattern which may provide a mechanism for regulating the expression or stability of either transcript (Ju et al., 2019; Vargas-Blanco & Shell, 2020).

The gene modules presented here are somewhat “blunt-force instruments’ applied to transcripts that are part of overlapping, coordinated responses to various environmental cues, but restricted to a single module grouping. Recent work exploring differentially expressed genes in response to various environmental conditions have revealed highly integrated adaptation responses. In other words, a single environmental change, e.g. hypoxia or growth on fatty acids or cholesterol, stimulates transcriptomic remodelling across diverse cellular functions, perhaps acting as cues to stimulate anticipatory pathways and ready the pathogen for the next challenge (Eoh et al., 2017; Gerrick et al., 2018). Confounders such as dual-function, ‘moonlighting’, proteins may weaken the correlation of a module with a specific condition and may create noise in otherwise well-connected modules. However, focussing on the best connected transcripts in various modules can uncover the unexpected connections between genes of diverse pathways.

Other methods of network analysis, such as those using deconvolution methods, allow genes to be members of more than one module and are considered less ‘noisy’ than clustering methods, such as WGCNA. However, these methods require extremely large numbers of samples to perform well, may be subject to batch effect issues between experimental datasets and characterise a limited proportion of the protein-coding transcripts expressed by Mtb (Saelens et al., 2018; Yoo, et al., 2022). Predicting ncRNA from different datasets involves a significant degree of quality control, parameter adjustment and manual curation, limiting the number of datasets that could be included in our analysis. Including more data would most likely strengthen the correlations with certain conditions and improve the overall specificity of the modules. However, the work presented here confirms that ncRNA are important players in adaptation responses, and their associations with the protein-coding genes in their assigned modules provides context for their activity.

The modules discussed in depth in this paper represent a limited snapshot of this extensive co-expression network. Modules of interest can be identified by correlations to experimental conditions, associated GO terms, functional categories, or gene group enrichment. The supplementary tables (Supp Table 2) have been organised into an easily-accessible spreadsheet for researchers to query particular genes or modules of interest and find associated protein-coding genes or ncRNA. These spreadsheets provide information about the module association, membership values, TSSs and for UTRs, the module membership of the adjacent ORFs for each predicted ncRNA. We anticipate this to be a useful resource for discovering ncRNA candidates for further investigation, add context to the circumstances of expression of previously identified ncRNAs, identify associations of genes with unknown functions and suggest roles for ‘moonlighting’ proteins that may be associated with unexpected gene groupings.

## Abbreviations

CDS: coding sequence
ME: module eigengene
MM: module membership
Mtb: *Mycobacterium tuberculosis*
MTBC: *Mycobacterium tuberculosis* complex
ncRNA: non-coding RNA
ORF: open reading frame
RNA-seq: RNA sequencing
RNAP: RNA polymerase
sORF: short open reading frame
sRNA: short non-coding RNA
TSS: transcription start site
TTS: transcription termination site
UTR: untranslated region
WGCNA: weighted gene co-expression analysis

## References

Aguilar-Ayala, D. A., Tilleman, L., Van Nieuwerburgh, F., Deforce, D., Palomino, J. C., Vandamme, P., Gonzalez-Y-Merchand, J. A., & Martin, A. (2017). The transcriptome of Mycobacterium tuberculosis in a lipid-rich dormancy model through RNAseq analysis. Scientific Reports, 7(1), 17665. https://doi.org/10.1038/s41598-017-17751-x

Ami, V. K. G., Balasubramanian, R., & Hegde, S. R. (2020). Genome-wide identification of the context-dependent sRNA expression in Mycobacterium tuberculosis. BMC Genomics, 21(167), 1–12.

Arnvig, K. B., Comas, I., Thomson, N. R., Houghton, J., Boshoff, H. I., Croucher, N. J., Rose, G., Perkins, T. T., Parkhill, J., Dougan, G., & Young, D. B. (2011). Sequence-Based Analysis Uncovers an Abundance of Non-Coding RNA in the Total Transcriptome of Mycobacterium tuberculosis. PLOS Pathogens, 7(11), e1002342. https://doi.org/10.1371/journal.ppat.1002342

Arnvig, K. B., & Young, D. B. (2009). Identification of small RNAs in Mycobacterium tuberculosis. Molecular Microbiology, 73(3), 397–408. https://doi.org/10.1111/j.1365-2958.2009.06777.x

Arnvig, K., & Young, D. (2012). Non-coding RNA and its potential role in Mycobacterium tuberculosis pathogenesis. RNA Biology, 9(4), 427–436. https://doi.org/10.4161/rna.20105

Ashburner, M., Ball, C. A., Blake, J. A., Botstein, D., Butler, H., Cherry, J. M., Davis, A. P., Dolinski, K., Dwight, S. S., Eppig, J. T., Harris, M. A., Hill, D. P., Issel-Tarver, L., Kasarskis, A., Lewis, S., Matese, J. C., Richardson, J. E., Ringwald, M., Rubin, G. M., & Sherlock, G. (2000). Gene Ontology: Tool for the unification of biology. Nature Genetics, 25(1), 25–29. https://doi.org/10.1038/75556

Becq, J., Gutierrez, M. C., Rosas-Magallanes, V., Rauzier, J., Gicquel, B., Neyrolles, O., & Deschavanne, P. (2007). Contribution of horizontally acquired genomic islands to the evolution of the tubercle bacilli. Molecular Biology and Evolution, 24(8), 1861–1871. https://doi.org/10.1093/molbev/msm111

Benjamini, Y., & Hochberg, Y. (1995). Controlling the False Discovery Rate: A Practical and Powerful Approach to Multiple Testing. Journal of the Royal Statistical Society: Series B (Methodological), 57(1), 289–300. https://doi.org/10.1111/j.2517-6161.1995.tb02031.x

Bhusal, R. P., Bashiri, G., Kwai, B. X. C., Sperry, J., & Leung, I. K. H. (2017). Targeting isocitrate lyase for the treatment of latent tuberculosis. Drug Discovery Today, 22(7), 1008–1016. https://doi.org/10.1016/j.drudis.2017.04.012

Bidnenko, E., & Bidnenko, V. (2018). Transcription termination factor Rho and microbial phenotypic heterogeneity. Current Genetics, 64(3), 541–546. https://doi.org/10.1007/s00294-017-0775-7

Blum, M., Chang, H.-Y., Chuguransky, S., Grego, T., Kandasaamy, S., Mitchell, A., Nuka, G., Paysan-Lafosse, T., Qureshi, M., Raj, S., Richardson, L., Salazar, G. A., Williams, L., Bork, P., Bridge, A., Gough, J., Haft, D. H., Letunic, I., Marchler-Bauer, A., … Finn, R. D. (2020). The InterPro protein families and domains database: 20 years on. Nucleic Acids Research, 49(D1), D344–D354. https://doi.org/10.1093/nar/gkaa977

Canestrari, J. G., Lasek-Nesselquist, E., Upadhyay, A., Rofaeil, M., Champion, M. M., Wade, J. T., Derbyshire, K. M., & Gray, T. A. (2020). Polycysteine-encoding leaderless short ORFs function as cysteine-responsive attenuators of operonic gene expression in mycobacteria. Molecular Microbiology, 114(1), 93–108. https://doi.org/10.1111/mmi.14498

Chao, Y., Papenfort, K., Reinhardt, R., Sharma, C. M., & Vogel, J. (2012). An atlas of Hfq-bound transcripts reveals 3’ UTRs as a genomic reservoir of regulatory small RNAs. The EMBO Journal, 31(20), 4005–4019. https://doi.org/10.1038/emboj.2012.229

Chen, J., & Xie, J. (2011). Role and regulation of bacterial LuxR-like regulators. Journal of Cellular Biochemistry, 112(10), 2694–2702. https://doi.org/10.1002/jcb.23219

Chetal, K., & Janga, S. C. (2015). OperomeDB: A Database of Condition-Specific Transcription Units in Prokaryotic Genomes. BioMed Research International, 2015, 318217–318217. PubMed. https://doi.org/10.1155/2015/318217

Cortes, T., Schubert, O. T., Rose, G., Arnvig, K. B., Comas, I., Aebersold, R., & Young, D. B. (2013). Genome-wide mapping of transcriptional start sites defines an extensive leaderless transcriptome in Mycobacterium tuberculosis. Cell Reports, 5(4), 1121– 1131. https://doi.org/10.1016/j.celrep.2013.10.031

Dar, D., Shamir, M., Mellin, J. R., Koutero, M., Stern-Ginossar, N., Cossart, P., & Sorek, R. (2016). Term-seq reveals abundant ribo-regulation of antibiotics resistance in bacteria. Science, 352(6282), aad9822. https://doi.org/10.1126/science.aad9822

Dar, D., & Sorek, R. (2018). Bacterial noncoding RNAs excised from within protein-coding transcripts. MBio, 9(5). https://doi.org/10.1128/mBio.01730-18

Del Portillo, P., García-Morales, L., Menéndez, M. C., Anzola, J. M., Rodríguez, J. G., Helguera-Repetto, A. C., Ares, M. A., Prados-Rosales, R., Gonzalez-y-Merchand, J. A., & García, M. J. (2019). Hypoxia Is Not a Main Stress When Mycobacterium tuberculosis Is in a Dormancy-Like Long-Chain Fatty Acid Environment. Frontiers in Cellular and Infection Microbiology, 8, 449–449.

Desgranges, E., Barrientos, L., & Caldelari, I. (2021). The 3’UTR-derived sRNA RsaG coordinates redox homeostasis and metabolism adaptation in response to glucose-6-phosphate uptake in Staphylococcus aureus. Molecular Microbiology. https://doi.org/10.1111/MMI.14845

D’Halluin, A., Polgar, P., Kipkorir, T., Patel, Z., Cortes, T., & Arnvig, K. B. (2022). Term-seq reveals an abundance of conditional, Rho-dependent termination in Mycobacterium tuberculosis. BioRxiv, 2022.06.01.494293. https://doi.org/10.1101/2022.06.01.494293

Dinan, Adam M., Tong, Pin, Lohan, Amanda J., Conlon, Kevin M., Miranda-CasoLuengo Aleksandra A., Malone, Kerri M., Gordon, Stephen V., & Loftus, Brendan J. (2014). Relaxed Selection Drives a Noisy Noncoding Transcriptome in Members of the Mycobacterium tuberculosis Complex. MBio, 5(4), e01169–14. https://doi.org/10.1128/mBio.01169-14

Du, P., Sohaskey, C. D., & Shi, L. (2016). Transcriptional and physiological changes during Mycobacterium tuberculosis reactivation from non-replicating persistence. Frontiers in Microbiology, 7(AUG). https://doi.org/10.3389/fmicb.2016.01346

Dutta, D. (2018). Advance in Research on Mycobacterium tuberculosis FabG4 and Its Inhibitor. Frontiers in Microbiology, 9. https://www.frontiersin.org/article/10.3389/fmicb.2018.01184

Ellis, M. J., & Haniford, D. B. (2016). Riboregulation of bacterial and archaeal transposition. WIREs RNA, 7(3), 382–398. https://doi.org/10.1002/wrna.1341

Eoh, H., Wang, Z., Layre, E., Rath, P., Morris, R., Branch Moody, D., & Rhee, K. Y. (2017). Metabolic anticipation in Mycobacterium tuberculosis. Nature Microbiology, 2(8), 17084. https://doi.org/10.1038/nmicrobiol.2017.84

Galperin, M. Y., Wolf, Y. I., Makarova, K. S., Vera Alvarez, R., Landsman, D., & Koonin, E. V. (2021). COG database update: Focus on microbial diversity, model organisms, and widespread pathogens. Nucleic Acids Research, 49(D1), D274–D281. https://doi.org/10.1093/nar/gkaa1018

Gerrick, E. R., Barbier, T., Chase, M. R., Xu, R., François, J., Lin, V. H., Szucs, M. J., Rock, J. M., Ahmad, R., Tjaden, B., Livny, J., & Fortune, S. M. (2018). Small RNA profiling in mycobacterium tuberculosis identifies mrsi as necessary for an anticipatory iron sparing response. Proceedings of the National Academy of Sciences of the United States of America, 115(25), 6464–6469. https://doi.org/10.1073/pnas.1718003115

Girardin, R. C., & McDonough, K. A. (2020). Small RNA Mcr11 requires the transcription factor AbmR for stable expression and regulates genes involved in the central metabolism of Mycobacterium tuberculosis. Molecular Microbiology, 113(2), 504–520. https://doi.org/10.1111/mmi.14436

Gonzalo-Asensio, J., Malaga, W., Pawlik, A., Astarie-Dequeker, C., Passemar, C., Moreau, F., Laval, F., Daffé, M., Martin, C., Brosch, R., & Guilhot, C. (2014). Evolutionary history of tuberculosis shaped by conserved mutations in the PhoPR virulence regulator. Proceedings of the National Academy of Sciences of the United States of America, 111(31), 11491–11496. https://doi.org/10.1073/pnas.1406693111

Gonzalo-Asensio, J., Mostowy, S., Harders-Westerveen, J., Huygen, K., Hernández-Pando, R., Thole, J., Behr, M., Gicquel, B., & Martín, C. (2008). PhoP: a missing piece in the intricate puzzle of Mycobacterium tuberculosis virulence. PloS One, 3(10), e3496–e3496. PubMed. https://doi.org/10.1371/journal.pone.0003496

Harold, L. K., Antoney, J., Ahmed, F. H., Hards, K., Carr, P. D., Rapson, T., Greening, C., Jackson, C. J., & Cook, G. M. (2019). FAD-sequestering proteins protect mycobacteria against hypoxic and oxidative stress. Journal of Biological Chemistry, 294(8), 2903–5814. https://doi.org/10.1074/jbc.RA118.006237

Houghton, Joanna, Rodgers, Angela, Rose, Graham, D’Halluin, Alexandre, Kipkorir, Terry, Barker, Declan, Waddell, Simon J., Arnvig, Kristine B., & Oglesby, Amanda G. (2021). The Mycobacterium tuberculosis sRNA F6 Modifies Expression of Essential Chaperonins, GroEL2 and GroES. Microbiology Spectrum, 9(2), e01095–21. https://doi.org/10.1128/Spectrum.01095-21

Huang, D. W., Sherman, B. T., & Lempicki, R. A. (2009a). Bioinformatics enrichment tools: Paths toward the comprehensive functional analysis of large gene lists. Nucleic Acids Research, 37(1), 1–13. https://doi.org/10.1093/nar/gkn923

Huang, D. W., Sherman, B. T., & Lempicki, R. A. (2009b). Systematic and integrative analysis of large gene lists using DAVID bioinformatics resources. Nature Protocols, 4(1), 44–57. https://doi.org/10.1038/nprot.2008.211

Ignatov, D. V., Salina, E. G., Fursov, M. V., Skvortsov, T. A., Azhikina, T. L., & Kaprelyants, A. S. (2015). Dormant non-culturable Mycobacterium tuberculosis retains stable low-abundant mRNA. BMC Genomics, 16(1), 954. https://doi.org/10.1186/s12864-015-2197-6

Jiang, J., Lin, C., Zhang, J., Wang, Y., Shen, L., Yang, K., Xiao, W., Li, Y., Zhang, L., & Liu, J. (2020). Transcriptome Changes of Mycobacterium marinum in the Process of Resuscitation From Hypoxia-Induced Dormancy. Frontiers in Genetics, 10(February), 1–13. https://doi.org/10.3389/fgene.2019.01359

Jiang, J., Sun, X., Wu, W., Li, L., Wu, H., Zhang, L., Yu, G., & Li, Y. (2016). Construction and application of a co-expression network in Mycobacterium tuberculosis. Scientific Reports, 6(March 2015), 1–18. https://doi.org/10.1038/srep28422

Jiao, X., Sherman, B. T., Huang, D. W., Stephens, R., Baseler, M. W., Lane, H. C., & Lempicki, R. A. (2012). DAVID-WS: a stateful web service to facilitate gene/protein list analysis. Bioinformatics, 28(13), 1805–1806. https://doi.org/10.1093/bioinformatics/bts251

Ju, X., Li, D., & Liu, S. (2019). Full-length RNA profiling reveals pervasive bidirectional transcription terminators in bacteria. Nature Microbiology, 4(11), 1907–1918. https://doi.org/10.1038/s41564-019-0500-z

Kanehisa, M., Sato, Y., & Kawashima, M. (2022). KEGG mapping tools for uncovering hidden features in biological data. Protein Science, 31(1), 47–53. https://doi.org/10.1002/pro.4172

Kapopoulou, A., Lew, J. M., & Cole, S. T. (2011). The MycoBrowser portal: A comprehensive and manually annotated resource for mycobacterial genomes. Tuberculosis, 91(1), 8–13. https://doi.org/10.1016/j.tube.2010.09.006

Kendall, S. L., Burgess, P., Balhana, R., Withers, M., Ten Bokum, A., Lott, J. S., Gao, C., Uhia-Castro, I., & Stoker, N. G. (2010). Cholesterol utilization in mycobacteria is controlled by two TetR-type transcriptional regulators: KstR and kstR2. Microbiology, 156(5), 1362–1371. https://doi.org/10.1099/mic.0.034538-0

Kendall, S. L., Withers, M., Soffair, C. N., Moreland, N. J., Gurcha, S., Sidders, B., Frita, R., Ten Bokum, A., Besra, G. S., Lott, J. S., & Stoker, N. G. (2007). A highly conserved transcriptional repressor controls a large regulon involved in lipid degradation in Mycobacterium smegmatis and Mycobacterium tuberculosis. Molecular Microbiology, 65(3), 684–699. https://doi.org/10.1111/j.1365-2958.2007.05827.x

Kipkorir, Terry, Mashabela, Gabriel T., de Wet, Timothy J., Koch, Anastasia, Dawes Stephanie S., Wiesner, Lubbe, Mizrahi, Valerie, Warner, Digby F., & Henkin, Tina M. (2021). De Novo Cobalamin Biosynthesis, Transport, and Assimilation and Cobalamin-Mediated Regulation of Methionine Biosynthesis in Mycobacterium smegmatis. Journal of Bacteriology, 203(7), e00620–20. https://doi.org/10.1128/JB.00620-20

Lamichhane, G., Arnvig, K. B., & McDonough, K. A. (2013). Definition and annotation of (myco)bacterial non-coding RNA. Tuberculosis, 93(1), 26–29. https://doi.org/10.1016/j.tube.2012.11.010

Langfelder, P., & Horvath, S. (2008). WGCNA: An R package for weighted correlation network analysis. BMC Bioinformatics, 9. https://doi.org/10.1186/1471-2105-9-559

Lejars, M., Kobayashi, A., & Hajnsdorf, E. (2019). Physiological roles of antisense RNAs in prokaryotes. Biochimie, 164, 3–16. https://doi.org/10.1016/j.biochi.2019.04.015

Li, Heng. (2013). *Aligning sequence reads, clone sequences and assembly contigs with BWA–MEM*. https://doi.org/10.48550/arXiv.1303.3997

Love, M. I., Huber, W., & Anders, S. (2014). Moderated estimation of fold change and dispersion for RNA-seq data with DESeq2. Genome Biology, 15(12), 1–21. https://doi.org/10.1186/s13059-014-0550-8

Lu, L., Wei, R., Bhakta, S., Waddell, S. J., & Boix, E. (2021). Weighted gene co-expression network analysis to identify key modules and hub genes associated with paucigranulocytic asthma. Antibiotics, 10(97). https://doi.org/10.3390/antibiotics10020097

Lunge, A., Gupta, R., Choudhary, E., & Agarwal, N. (2020). The unfoldase ClpC1 of Mycobacterium tuberculosis regulates the expression of a distinct subset of proteins having intrinsically disordered termini. Journal of Biological Chemistry, 295(28), 9455–9473. https://doi.org/10.1074/jbc.RA120.013456

Maciąg Anna, Dainese Elisa, Rodriguez G. Marcela, Milano Anna, Provvedi Roberta, Pasca Maria R., Smith Issar, Palù Giorgio, Riccardi Giovanna, & Manganelli Riccardo. (2007). Global Analysis of the Mycobacterium tuberculosis Zur (FurB) Regulon. Journal of Bacteriology, 189(3), 730–740. https://doi.org/10.1128/JB.01190-06

Mai, J., Rao, C., Watt, J., Sun, X., Lin, C., Zhang, L., & Liu, J. (2019). Mycobacterium tuberculosis 6C sRNA binds multiple mRNA targets via C-rich loops independent of RNA chaperones. Nucleic Acids Research, 47(8), 4292–4307. https://doi.org/10.1093/nar/gkz149

Martini, M. C., Zhou, Y., Sun, H., & Shell, S. S. (2019). Defining the Transcriptional and Post-transcriptional Landscapes of Mycobacterium smegmatis in Aerobic Growth and Hypoxia. In Frontiers in Microbiology (Vol. 10). https://www.frontiersin.org/article/10.3389/fmicb.2019.00591

Menendez-Gil, P., Caballero, C., Catalan-Moreno, A., Irurzun, N., Barrio-Hernandez, I., Caldelari, I., & Toledo-Arana, A. (2020). Differential evolution in 3’UTRs leads to specific gene expression in Staphylococcus. Nucleic Acids Research, 48. https://doi.org/10.1093/nar/gkaa047

Menendez-Gil, P., & Toledo-Arana, A. (2021). Bacterial 3’UTRs: A Useful Resource in Post-transcriptional Regulation. Frontiers in Molecular Biosciences, 7. https://www.frontiersin.org/article/10.3389/fmolb.2020.617633

Modlin, S. J., Afif, E., Deepika, G., Zlotnicki, A. M., Dillon, N. A., Dhillon, N., Kuo, N., Robinhold, C., Chan, C. K., Baughn, A. D., & Valafar, F. (2021). Structure-Aware Mycobacterium tuberculosis Functional Annotation Uncloaks Resistance, Metabolic, and Virulence Genes. MSystems, 0(0), e00673–21. https://doi.org/10.1128/mSystems.00673-21

Moores, A., Riesco, A. B., Schwenk, S., & Arnvig, K. B. (2017). Expression, maturation and turnover of DrrS, an unusually stable, DosR regulated small RNA in Mycobacterium tuberculosis. PLOS ONE, 12(3), e0174079. https://doi.org/10.1371/journal.pone.0174079

Nesbitt, N. M., Yang, X., Fontán, P., Kolesnikova, I., Smith, I., Sampson, N. S., & Dubnau, E. (2010). A Thiolase of Mycobacterium tuberculosis Is Required for Virulence and Production of Androstenedione and Androstadienedione from Cholesterol. Infection and Immunity, 78(1), 275 LP – 282. https://doi.org/10.1128/IAI.00893-09

Ozuna, A., Liberto, D., Joyce, R. M., Arnvig, K. B., & Nobeli, I. (2019). baerhunter: An R package for the discovery and analysis of expressed non-coding regions in bacterial RNA-seq data. Bioinformatics. https://doi.org/10.1093/bioinformatics/btz643

Pawełczyk, J., Brzostek, A., Minias, A., Płociński, P., Rumijowska-Galewicz, A., Strapagiel, D., Zakrzewska-Czerwińska, J., & Dziadek, J. (2021). Cholesterol-dependent transcriptome remodeling reveals new insight into the contribution of cholesterol to Mycobacterium tuberculosis pathogenesis. Scientific Reports, 11(1), 12396. https://doi.org/10.1038/s41598-021-91812-0

Ponath, F., Hör, J., & Vogel, J. (2022). An overview of gene regulation in bacteria by small RNAs derived from mRNA 3’ ends. FEMS Microbiology Reviews, fuac017. https://doi.org/10.1093/femsre/fuac017

Puniya, B. L., Kulshreshtha, D., Verma, S. P., Kumar, S., & Ramachandran, S. (2013). Integrated gene co-expression network analysis in the growth phase of Mycobacterium tuberculosis reveals new potential drug targets. Molecular BioSystems, 9(11), 2798–2815. https://doi.org/10.1039/c3mb70278b

Ramage, H. R., Connolly, L. E., & Cox, J. S. (2009). Comprehensive functional analysis of Mycobacterium tuberculosis toxin-antitoxin systems: Implications for pathogenesis, stress responses, and evolution. PLoS Genetics, 5(12). https://doi.org/10.1371/journal.pgen.1000767

Ritchie, M. E., Phipson, B., Wu, D., Hu, Y., Law, C. W., Shi, W., & Smyth, G. K. (2015). Limma powers differential expression analyses for RNA-sequencing and microarray studies. Nucleic Acids Research, 43(7), e47–e47. https://doi.org/10.1093/nar/gkv007

Rustad, T. R., Harrell, M. I., Liao, R., & Sherman, D. R. (2008). The enduring hypoxic response of Mycobacterium tuberculosis. PLoS ONE, 3(1), 1–8. https://doi.org/10.1371/journal.pone.0001502

Saelens, W., Cannoodt, R., & Saeys, Y. (2018). A comprehensive evaluation of module detection methods for gene expression data. Nature Communications, 9(1), 1090. https://doi.org/10.1038/s41467-018-03424-4

Sáenz-Lahoya S., Bitarte N., García B., Burgui S., Vergara-Irigaray M., Valle J., Solano C., Toledo-Arana A., & Lasa I. (2019). Noncontiguous operon is a genetic organization for coordinating bacterial gene expression. Proceedings of the National Academy of Sciences, 116(5), 1733–1738. https://doi.org/10.1073/pnas.1812746116

Sala, A., Bordes, P., & Genevaux, P. (2014). Multiple toxin-antitoxin systems in Mycobacterium tuberculosis. Toxins, 6(3), 1002–1020. https://doi.org/10.3390/toxins6031002

Sawyer, E. B., Phelan, J. E., Clark, T. G., & Cortes, T. (2021). A snapshot of translation in Mycobacterium tuberculosis during exponential growth and nutrient starvation revealed by ribosome profiling. Cell Reports, 34(5). https://doi.org/10.1016/j.celrep.2021.108695

Schwenk, S., & Arnvig, K. B. (2018). Regulatory RNA in Mycobacterium tuberculosis, back to basics. Pathogens and Disease, 76(4). https://doi.org/10.1093/femspd/fty035

Serafini, A., Pisu, D., Palù, G., Rodriguez, G. M., & Manganelli, R. (2013). The ESX-3 Secretion System Is Necessary for Iron and Zinc Homeostasis in Mycobacterium tuberculosis. PLoS ONE, 8(10), 1–15. https://doi.org/10.1371/journal.pone.0078351

Shell, S. S., Wang, J., Lapierre, P., Mir, M., Chase, M. R., Pyle, M. M., Gawande, R., Ahmad, R., Sarracino, D. A., Ioerger, T. R., Fortune, S. M., Derbyshire, K. M., Wade, J. T., & Gray, T. A. (2015). Leaderless Transcripts and Small Proteins Are Common Features of the Mycobacterial Translational Landscape. PLOS Genetics, 11(11), e1005641. https://doi.org/10.1371/journal.pgen.1005641

Shockey, A. C., Dabney, J., & Pepperell, C. S. (2019). Effects of Host, Sample, and in vitro Culture on Genomic Diversity of Pathogenic Mycobacteria. In Frontiers in Genetics (Vol. 10). https://www.frontiersin.org/article/10.3389/fgene.2019.00477

Šiková, M., Janoušková, M., Ramaniuk, O., Páleníková, P., Pospíšil, J., Bartl, P., Suder, A., Pajer, P., Kubičková, P., Pavliš, O., Hradilová, M., Vítovská, D., Šanderová, H., Převorovský, M., Hnilicová, J., & Krásný, L. (2019). Ms1 RNA increases the amount of RNA polymerase in Mycobacterium smegmatis. Molecular Microbiology, 111(2), 354–372. https://doi.org/10.1111/mmi.14159

Singh Prabhat Ranjan, Vijjamarri Anil Kumar, Sarkar Dibyendu, & Federle Michael J. (2020). Metabolic Switching of Mycobacterium tuberculosis during Hypoxia Is Controlled by the Virulence Regulator PhoP. Journal of Bacteriology, 202(7), e00705–19. https://doi.org/10.1128/JB.00705-19

Smith, C., Canestrari, J. G., Wang, A. J., Champion, M. M., Derbyshire, K. M., Gray, T. A., & Wade, J. T. (2022). Pervasive translation in Mycobacterium tuberculosis. ELife, 11. e73980. https://doi.org/10.7554/eLife.73980

Solans, L., Gonzalo-Asensio, J., Sala, C., Benjak, A., Uplekar, S., Rougemont, J., Guilhot, C., Malaga, W., Martín, C., & Cole, S. T. (2014). The PhoP-Dependent ncRNA Mcr7 Modulates the TAT Secretion System in Mycobacterium tuberculosis. PLOS Pathogens, 10(5), e1004183. https://doi.org/10.1371/journal.ppat.1004183

Sreenu, V. B., Kumar, P., Nagaraju, J., & Nagarajaram, H. A. (2007). Simple sequence repeats in mycobacterial genomes. Journal of Biosciences, 32(1), 3–15. https://doi.org/10.1007/s12038-007-0002-7

Stiens, J., Arnvig, K. B., Kendall, S. L., & Nobeli, I. (2022). Challenges in defining the functional, non-coding, expressed genome of members of the Mycobacterium tuberculosis complex. Molecular Microbiology, 117(1), 20–31. https://doi.org/10.1111/mmi.14862

Talwar, S., Pandey, M., Sharma, C., Kutum, R., Lum, J., Carbajo, D., Goel, R., Poidinger, M., Dash, D., Singhal, A., & Pandey, A. K. (2020). Role of VapBC12 Toxin-Antitoxin Locus in Cholesterol-Induced Mycobacterial Persistence. MSystems, 5(6). https://doi.org/10.1128/msystems.00855-20

The Gene Ontology Consortium. (2021). The Gene Ontology resource: Enriching a GOld mine. Nucleic Acids Research, 49(D1), D325–D334. https://doi.org/10.1093/nar/gkaa1113

Toledo-Arana, A., & Lasa, I. (2020). Advances in bacterial transcriptome understanding: From overlapping transcription to the excludon concept. Molecular Microbiology, 113(3), 593–602. https://doi.org/10.1111/mmi.14456

Updegrove, T. B., Kouse, A. B., Bandyra, K. J., & Storz, G. (2019). Stem-loops direct precise processing of 3’ UTR-derived small RNA MicL. Nucleic Acids Research, 47(3), 1482–1492. https://doi.org/10.1093/nar/gky1175

Vargas-Blanco, D. A., & Shell, S. S. (2020). Regulation of mRNA Stability During Bacterial Stress Responses. Frontiers in Microbiology, 11(September). https://doi.org/10.3389/fmicb.2020.02111

Voskuil, M. I., Visconti, K. C., & Schoolnik, G. K. (2004). Mycobacterium tuberculosis gene expression during adaptation to stationary phase and low-oxygen dormancy. Tuberculosis, 84(3–4), 218–227. https://doi.org/10.1016/j.tube.2004.02.003

Wade, J. T., & Grainger, D. C. (2014). Pervasive transcription: Illuminating the dark matter of bacterial transcriptomes. Nature Reviews Microbiology, 12(9), 647–653. https://doi.org/10.1038/nrmicro3316

Wang, X., Monford Paul Abishek, N., Jeon, H. J., Lee, Y., He, J., Adhya, S., & Lim, H. M. (2019). Processing generates 3’ ends of RNA masking transcription termination events in prokaryotes. Proceedings of the National Academy of Sciences of the United States of America, 116(10), 4440–4445. https://doi.org/10.1073/pnas.1813181116

Warman, E. A., Forrest, D., Guest, T., Haycocks, J. J. R. J., Wade, J. T., & Grainger, D. C. (2021). Widespread divergent transcription from bacterial and archaeal promoters is a consequence of DNA-sequence symmetry. Nature Microbiology, 6(6), 746–756. https://doi.org/10.1038/s41564-021-00898-9

Warner, D. F., Savvi, S., Mizrahi, V., & Dawes, S. S. (2007). A Riboswitch Regulates Expression of the Coenzyme B12-Independent Methionine Synthase in Mycobacterium tuberculosis: Implications for Differential Methionine Synthase Function in Strains H37Rv and CDC1551. Journal of Bacteriology, 189(9), 3655 LP – 3659. https://doi.org/10.1128/JB.00040-07

World Health Organization. (2021, October 14). Tuberculosis Fact Sheet. Tuberculosis. https://www.who.int/news-room/fact-sheets/detail/tuberculosis

Yoo, Reo, Rychel, Kevin, Poudel, Saugat, Al-bulushi, Tahani, Yuan Yuan, Chauhan, Siddharth, Lamoureux, Cameron, Palsson, Bernhard O., Sastry, Anand, & Tringe, Susannah Green. (2022). Machine Learning of All Mycobacterium tuberculosis H37Rv RNA-seq Data Reveals a Structured Interplay between Metabolism, Stress Response, and Infection. MSphere, 0(0), e00033–22. https://doi.org/10.1128/msphere.00033-22

Zhang, B., & Horvath, S. (2005). A General Framework for Weighted Gene Co-Expression Network Analysis. Statistical Applications in Genetics and Molecular Biology, 4(1). https://doi.org/10.2202/1544-6115.1128

Zhang, L., Hendrickson, R. C., Meikle, V., Lefkowitz, E. J., Ioerger, T. R., & Niederweis, M. (2020). Comprehensive analysis of iron utilization by Mycobacterium tuberculosis. PLOS Pathogens, 16(2), e1008337. https://doi.org/10.1371/journal.ppat.1008337

Zhou, Y., Huang, H., Zhou, P., & Xie, J. (2012). Molecular mechanisms underlying the function diversity of transcriptional factor IclR family. Cellular Signalling, 24(6), 1270–1275. https://doi.org/10.1016/j.cellsig.2012.02.008

Zondervan, N. A., Van Dam, J. C. J., Schaap, P. J., Martins dos Santos, V. A. P., & Suarez-Diez, M. (2018). Regulation of Three Virulence Strategies of Mycobacterium tuberculosis: A Success Story. International Journal of Molecular Sciences, 19(2). https://doi.org/10.3390/ijms19020347

