## Supplemental Figures for "Using a Whole Genome Co-expression Network to Inform the Functional Characterisation of Predicted Genomic Elements from *Mycobacterium tuberculosis* Transcriptomic Data"

### Slide 1
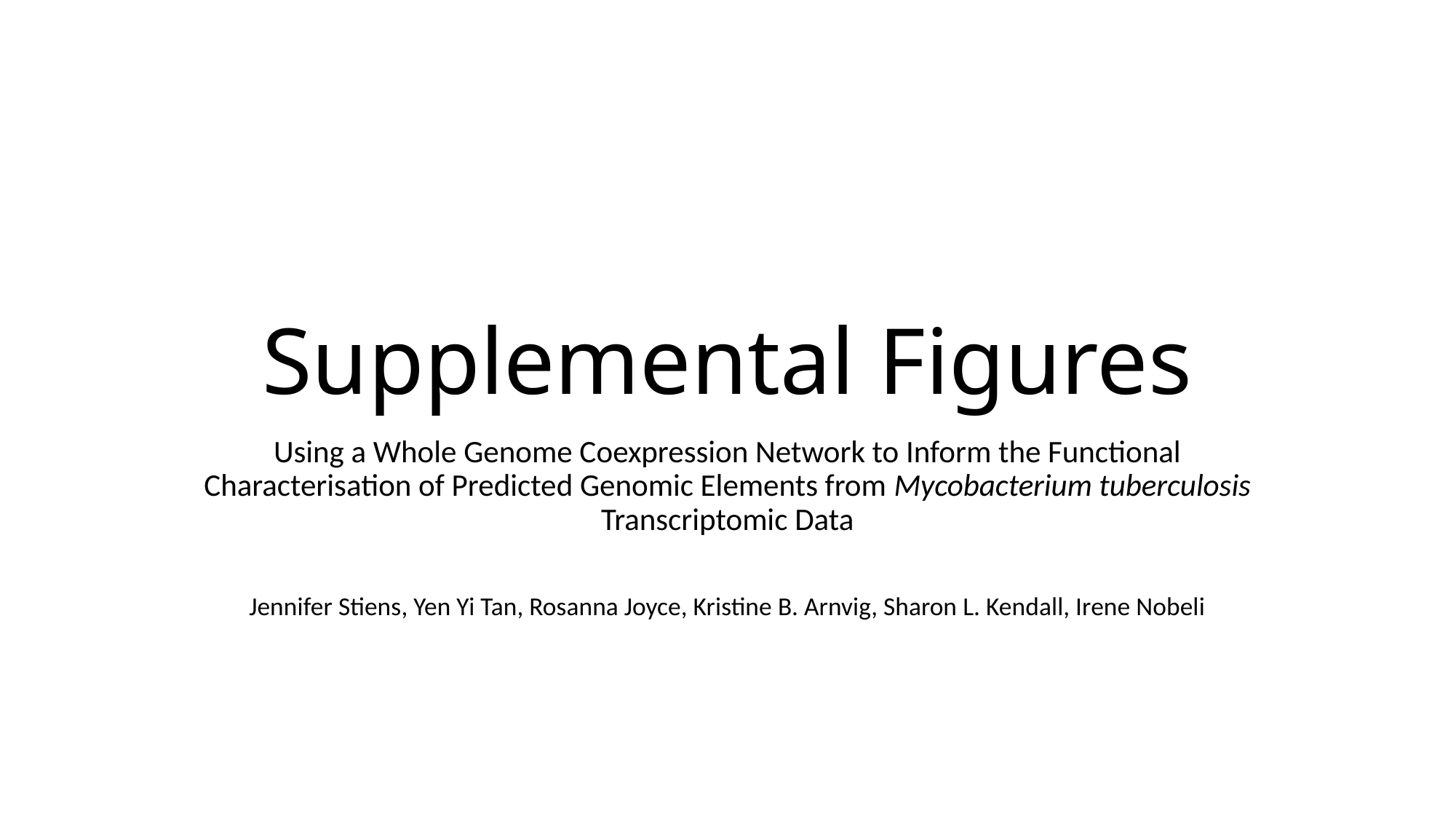

#### Supplemental Figures
Using a Whole Genome Coexpression Network to Inform the Functional Characterisation of Predicted Genomic Elements from Mycobacterium tuberculosis Transcriptomic Data
Jennifer Stiens, Yen Yi Tan, Rosanna Joyce, Kristine B. Arnvig, Sharon L. Kendall, Irene Nobeli

### Slide 2
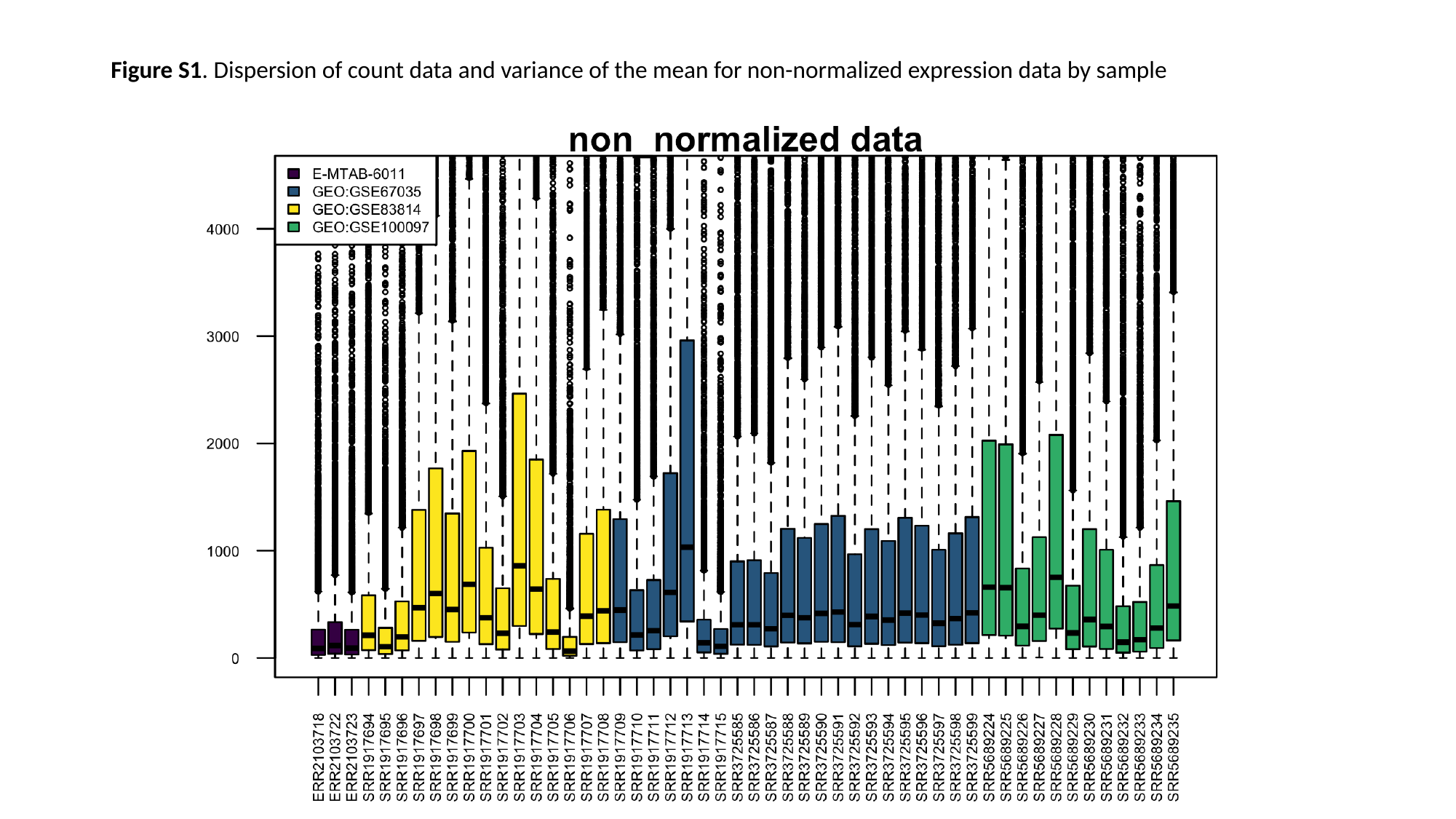

#### Figure S1. Dispersion of count data and variance of the mean for non-normalized expression data by sample

### Slide 3
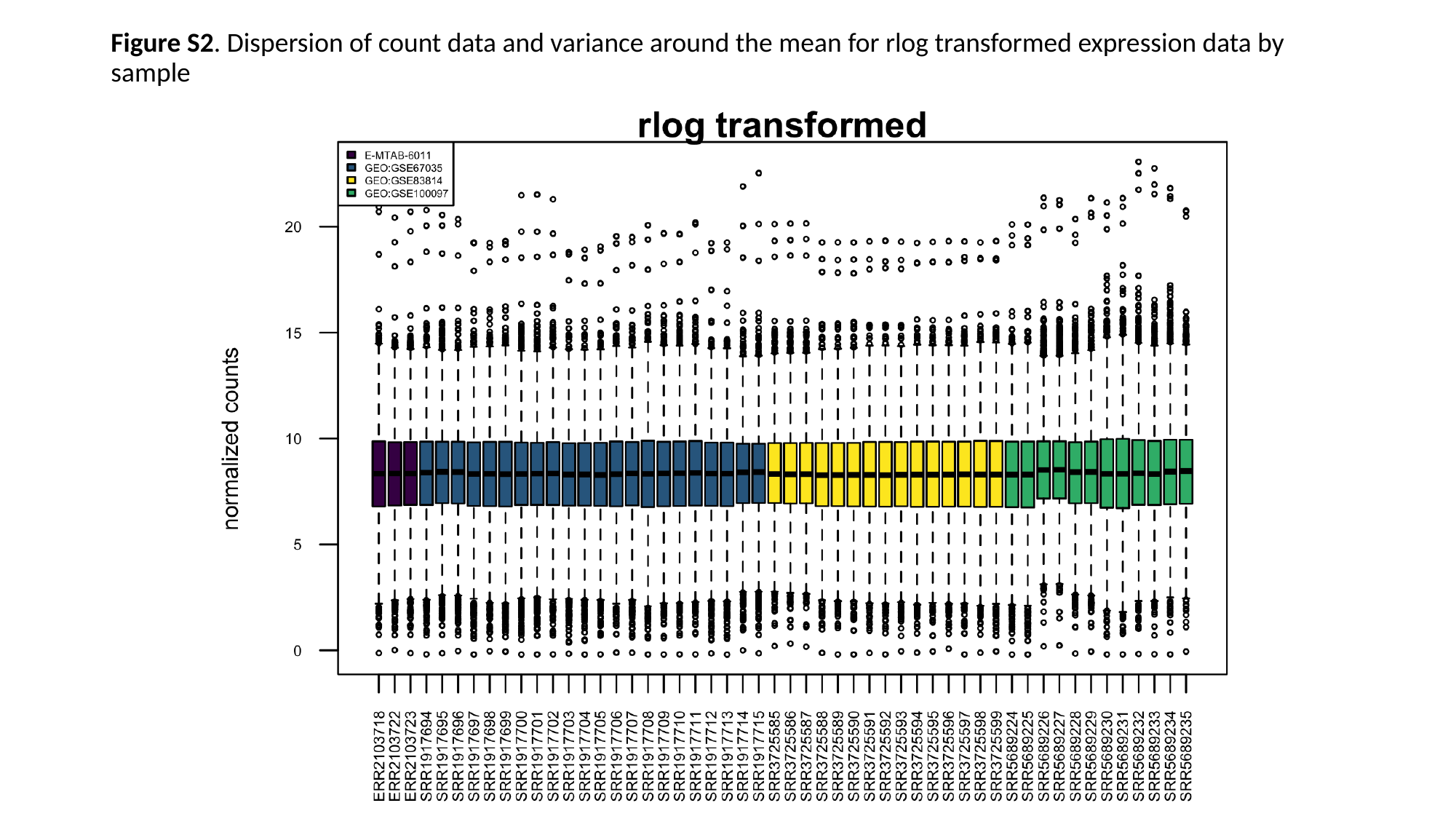

#### Figure S2. Dispersion of count data and variance around the mean for rlog transformed expression data by sample

### Slide 4
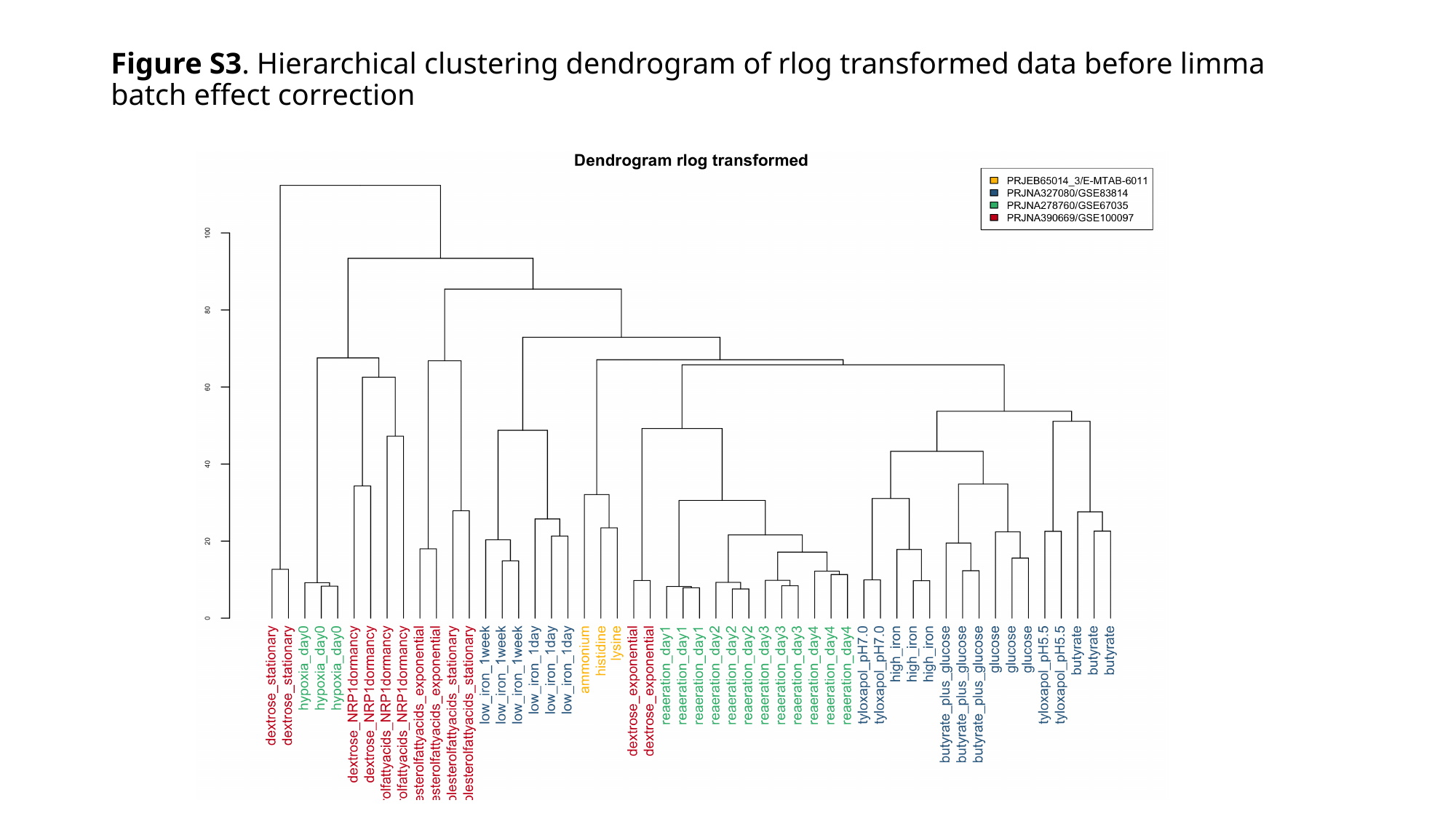

#### Figure S3. Hierarchical clustering dendrogram of rlog transformed data before limma batch effect correction

### Slide 5
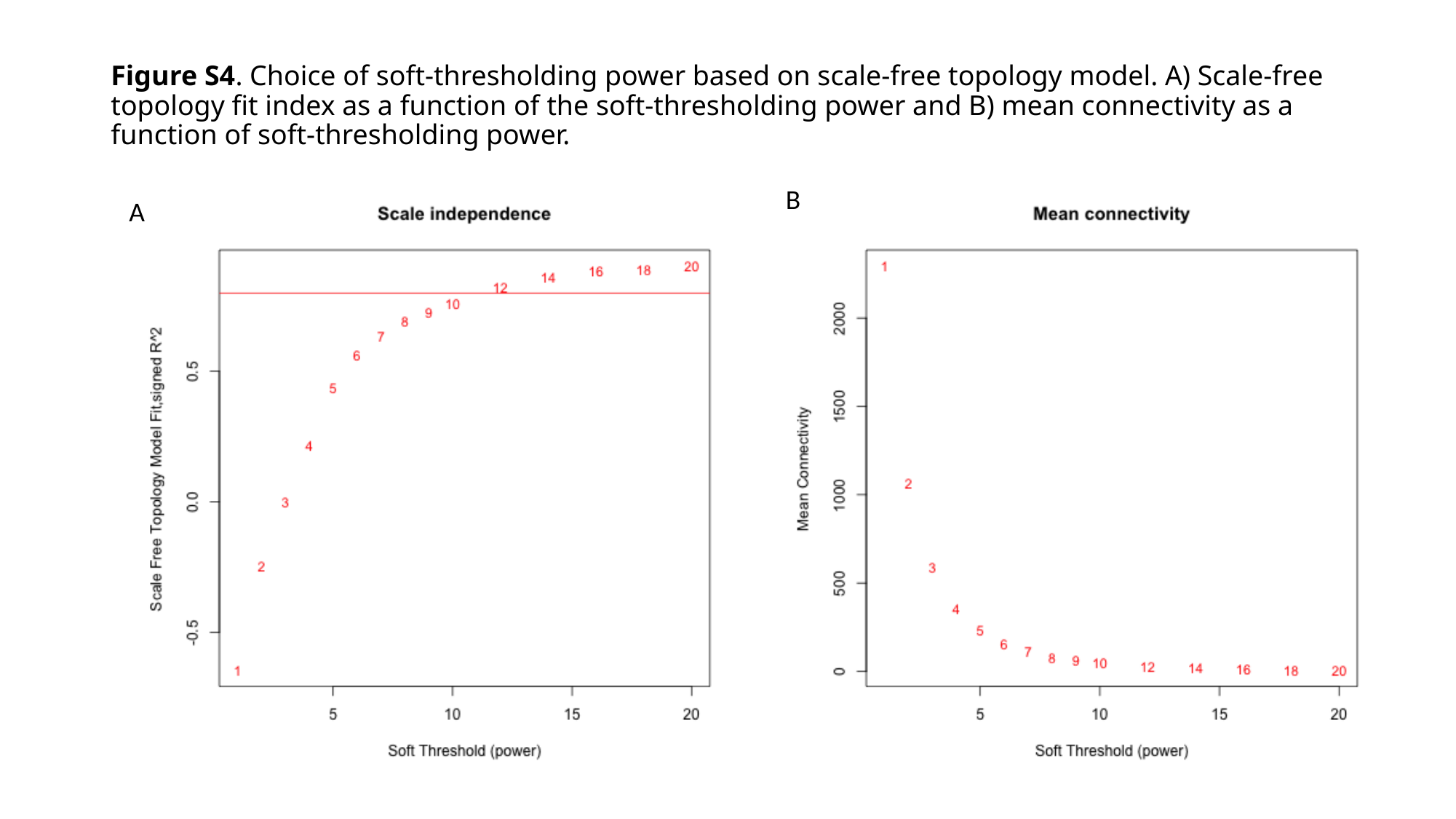

#### Figure S4. Choice of soft-thresholding power based on scale-free topology model. A) Scale-free topology fit index as a function of the soft-thresholding power and B) mean connectivity as a function of soft-thresholding power.
B
A

### Slide 6
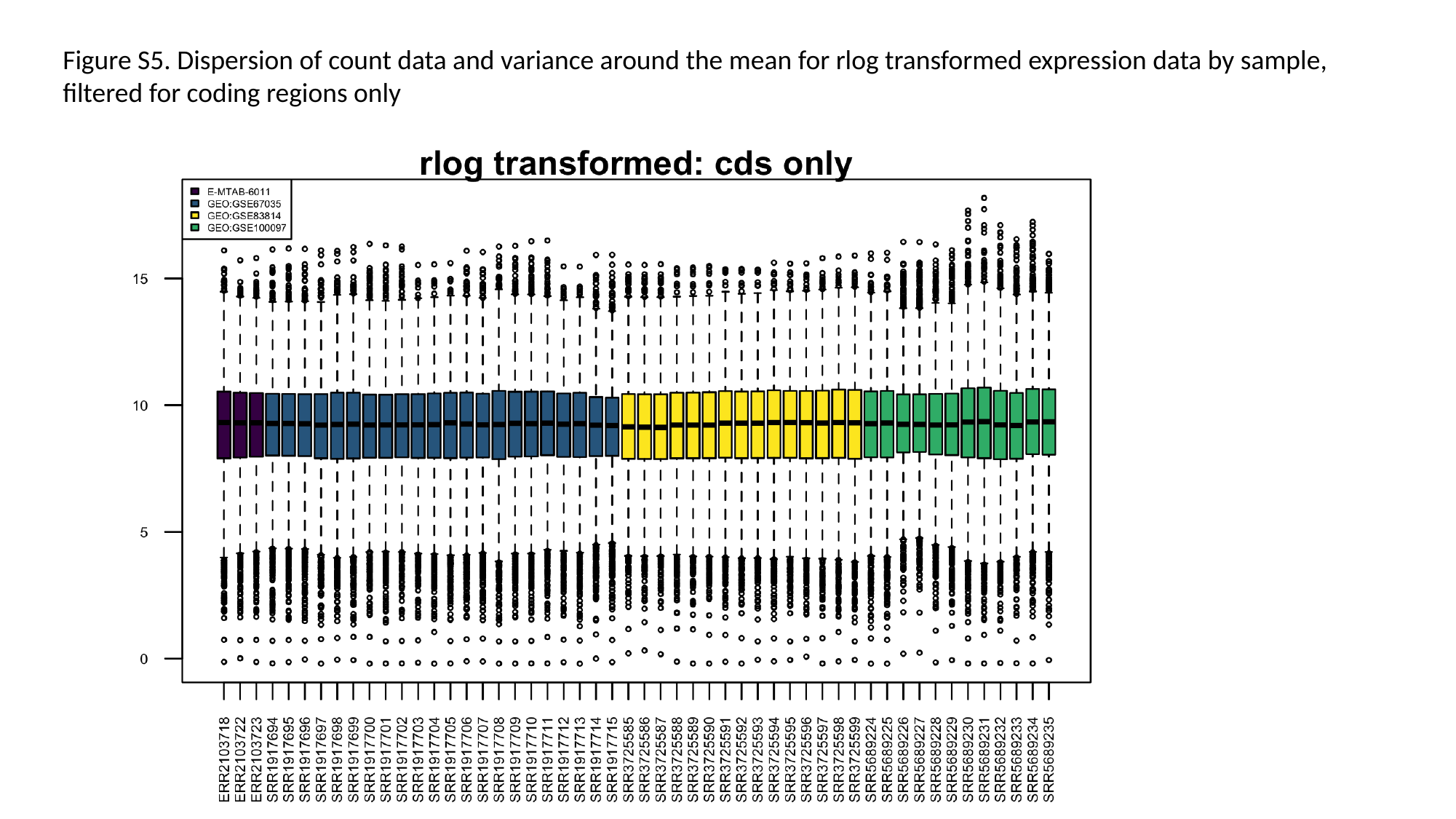

Figure S5. Dispersion of count data and variance around the mean for rlog transformed expression data by sample, filtered for coding regions only

### Slide 7
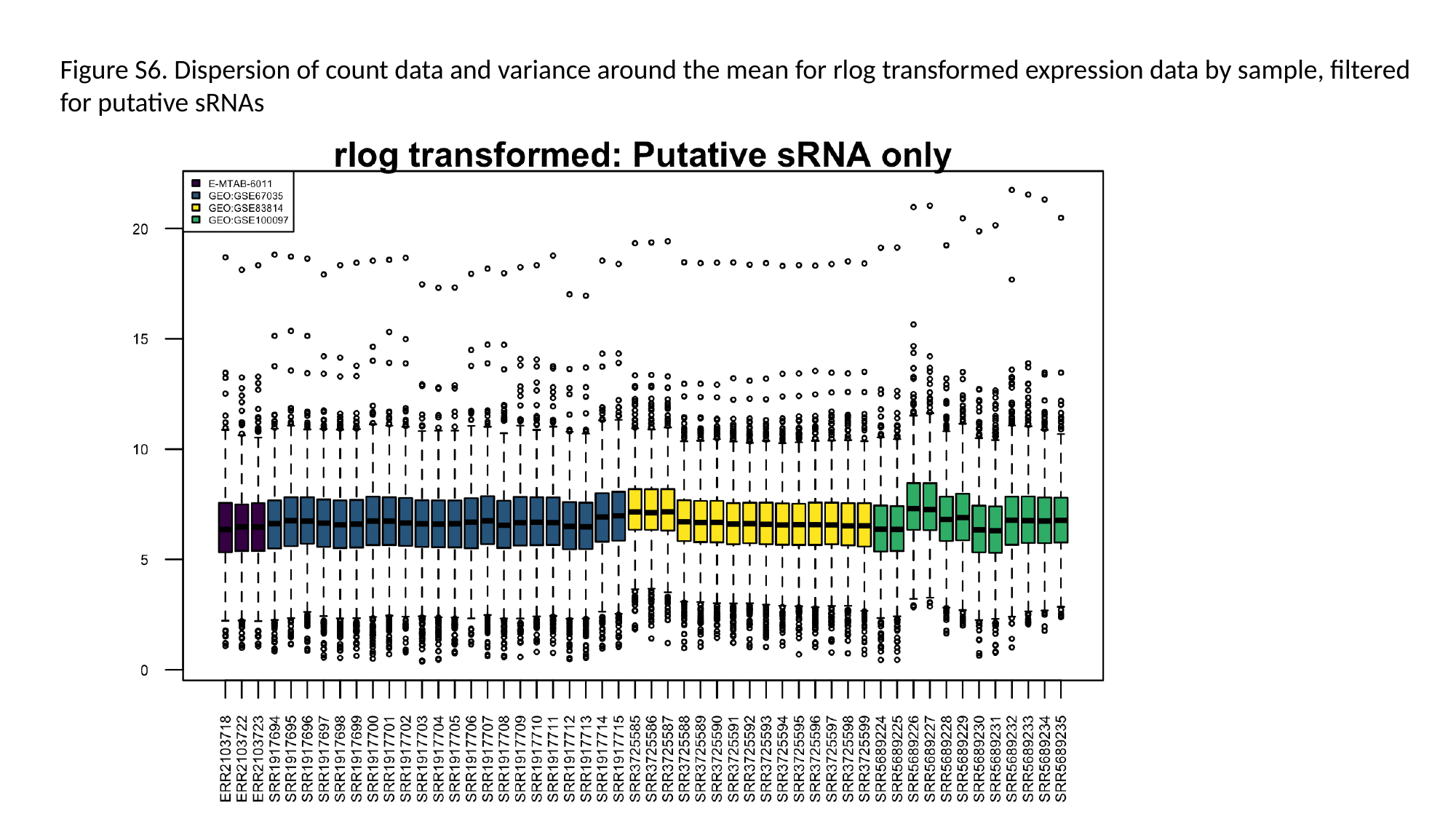

Figure S6. Dispersion of count data and variance around the mean for rlog transformed expression data by sample, filtered for putative sRNAs

### Slide 8
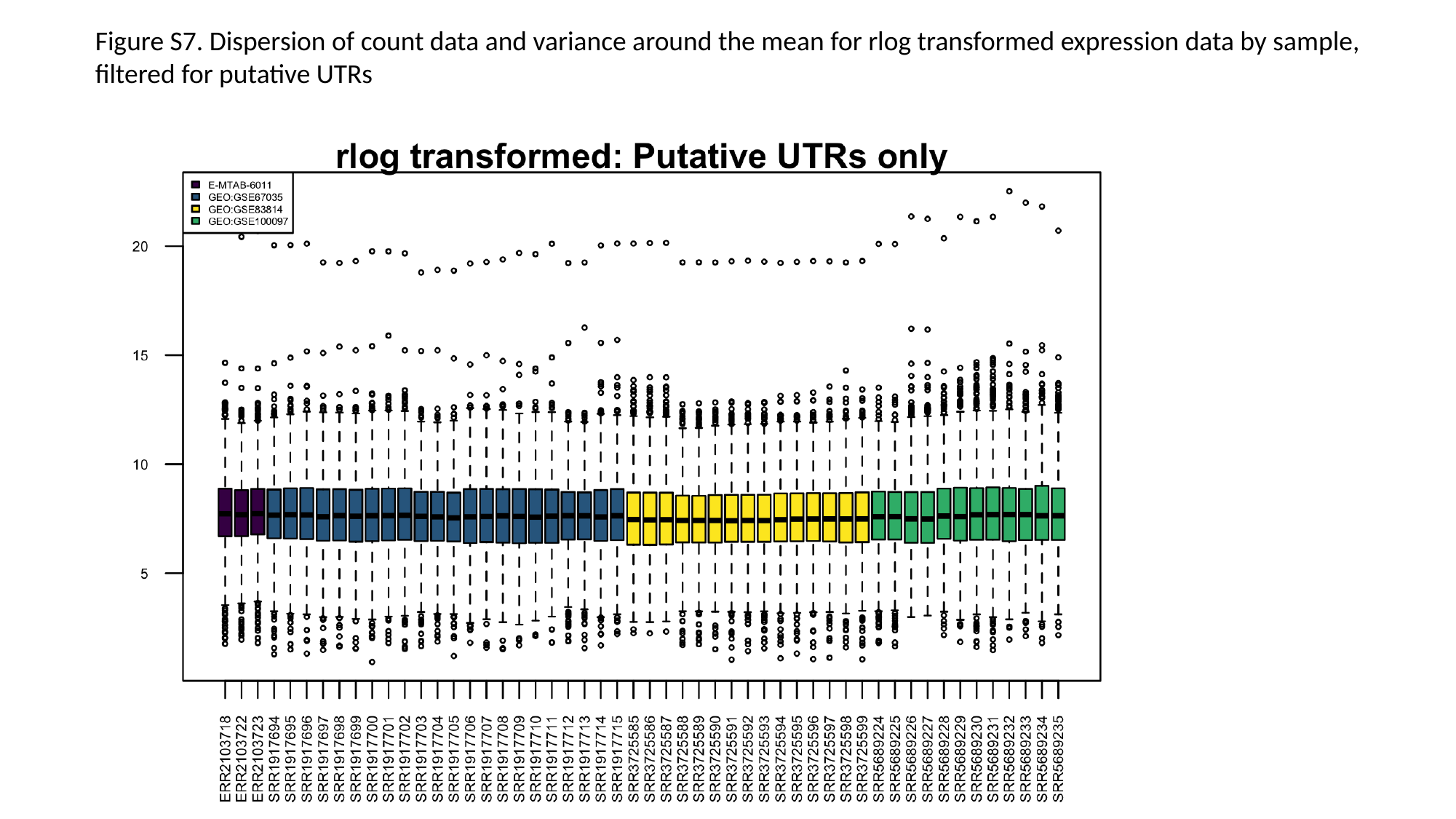

Figure S7. Dispersion of count data and variance around the mean for rlog transformed expression data by sample, filtered for putative UTRs

### Slide 9
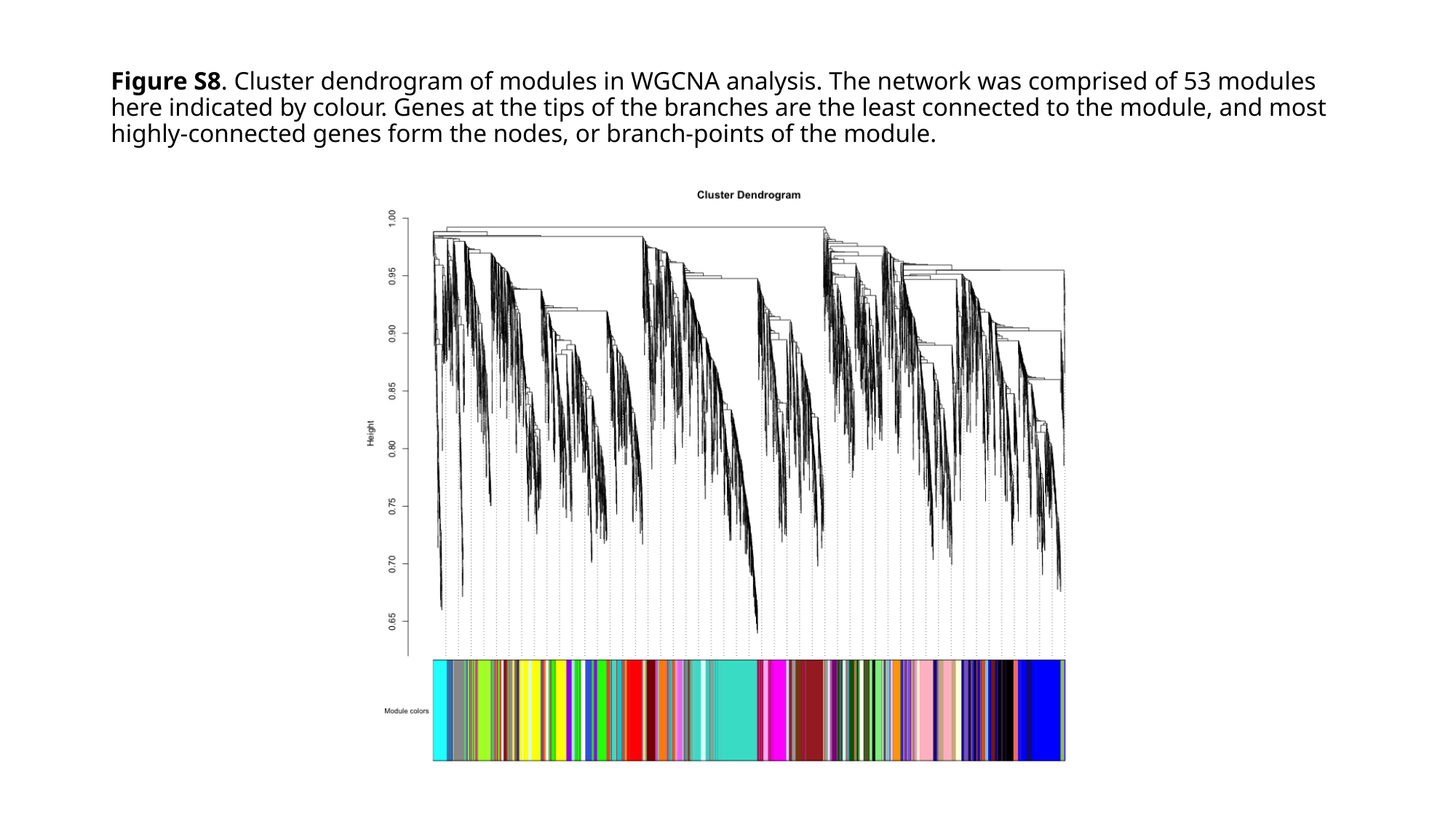

#### Figure S8. Cluster dendrogram of modules in WGCNA analysis. The network was comprised of 53 modules here indicated by colour. Genes at the tips of the branches are the least connected to the module, and most highly-connected genes form the nodes, or branch-points of the module.

### Slide 10
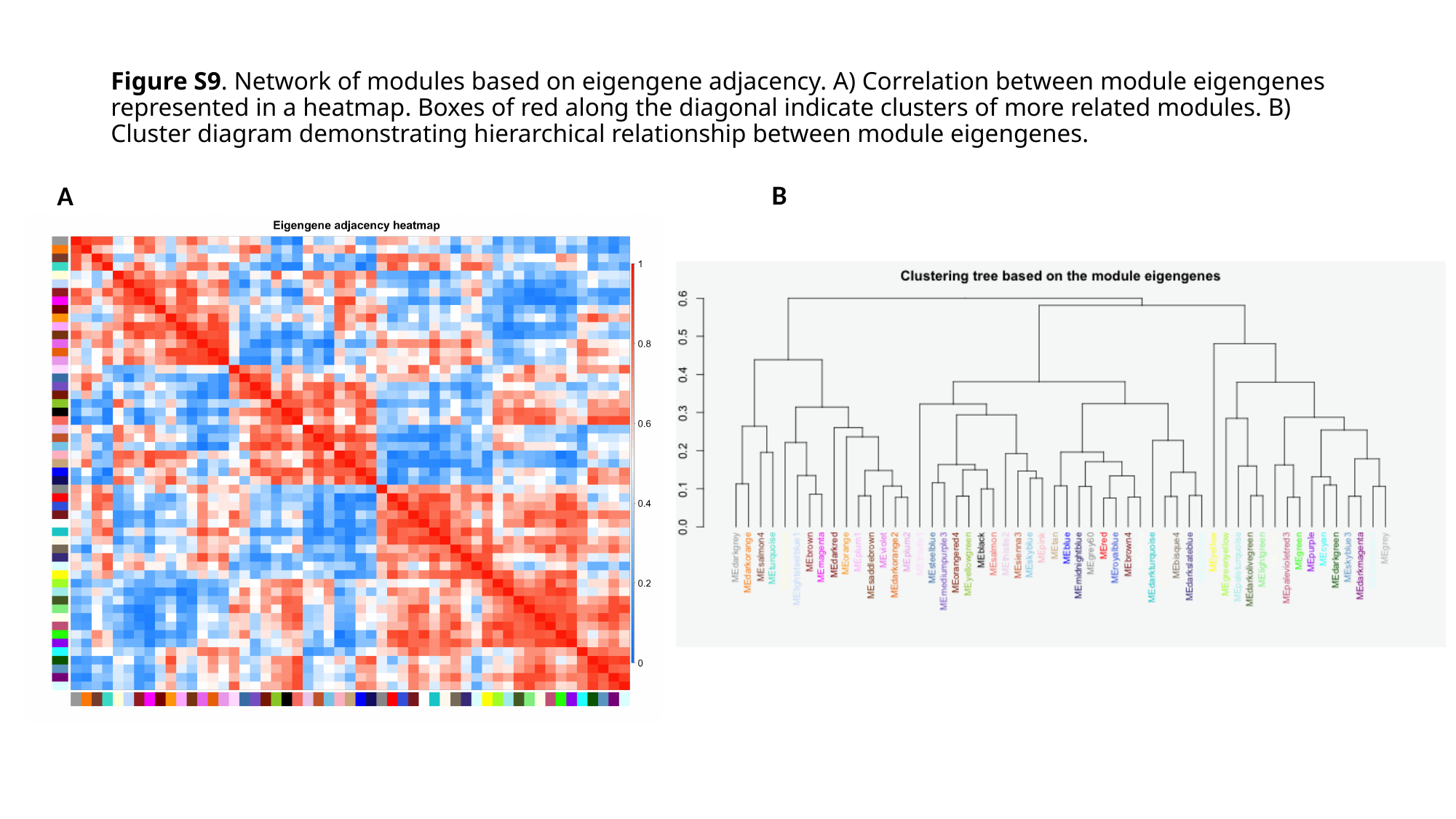

#### Figure S9. Network of modules based on eigengene adjacency. A) Correlation between module eigengenes represented in a heatmap. Boxes of red along the diagonal indicate clusters of more related modules. B) Cluster diagram demonstrating hierarchical relationship between module eigengenes.
B
A

### Slide 11
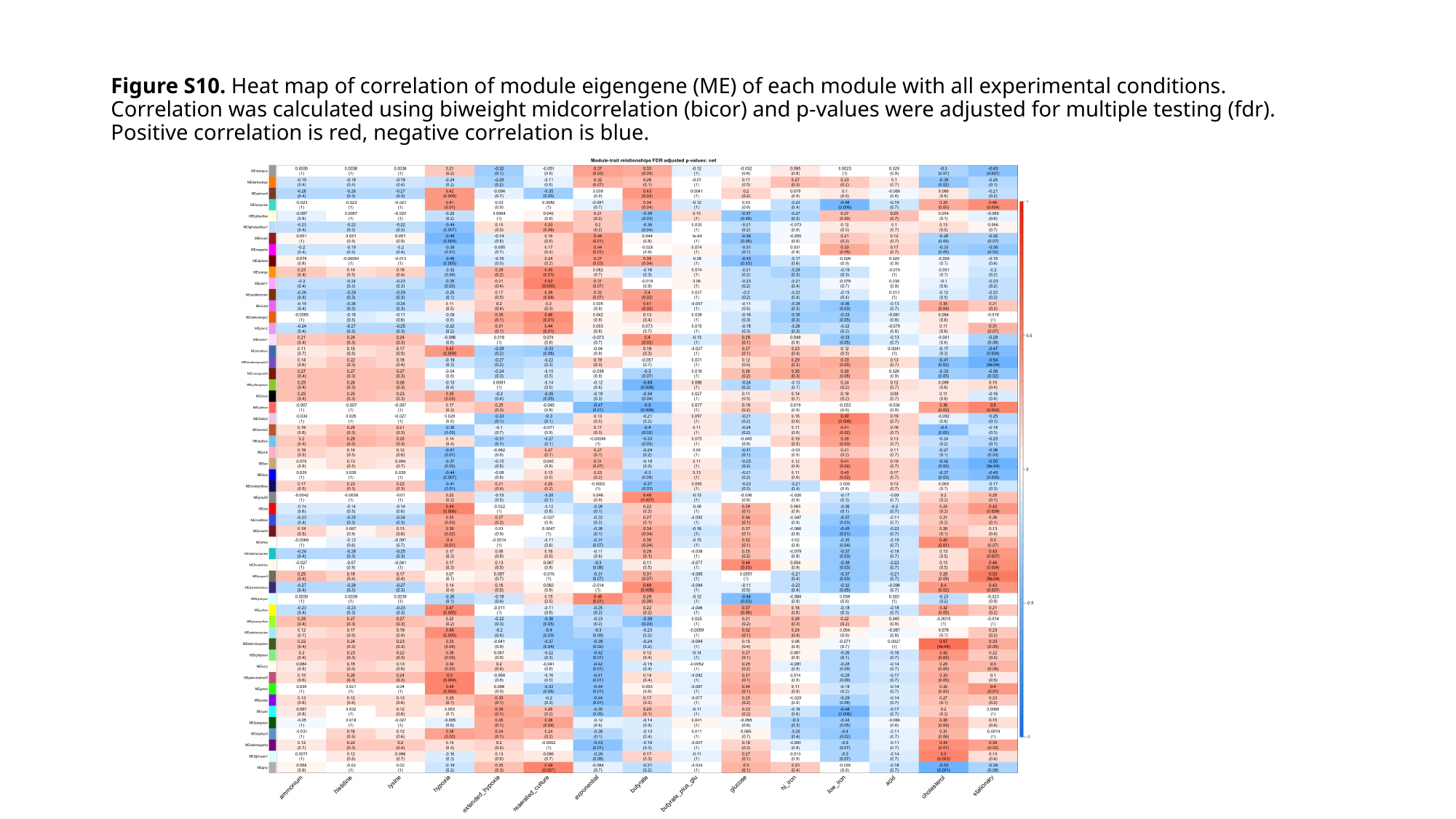

#### Figure S10. Heat map of correlation of module eigengene (ME) of each module with all experimental conditions. Correlation was calculated using biweight midcorrelation (bicor) and p-values were adjusted for multiple testing (fdr). Positive correlation is red, negative correlation is blue.

### Slide 12
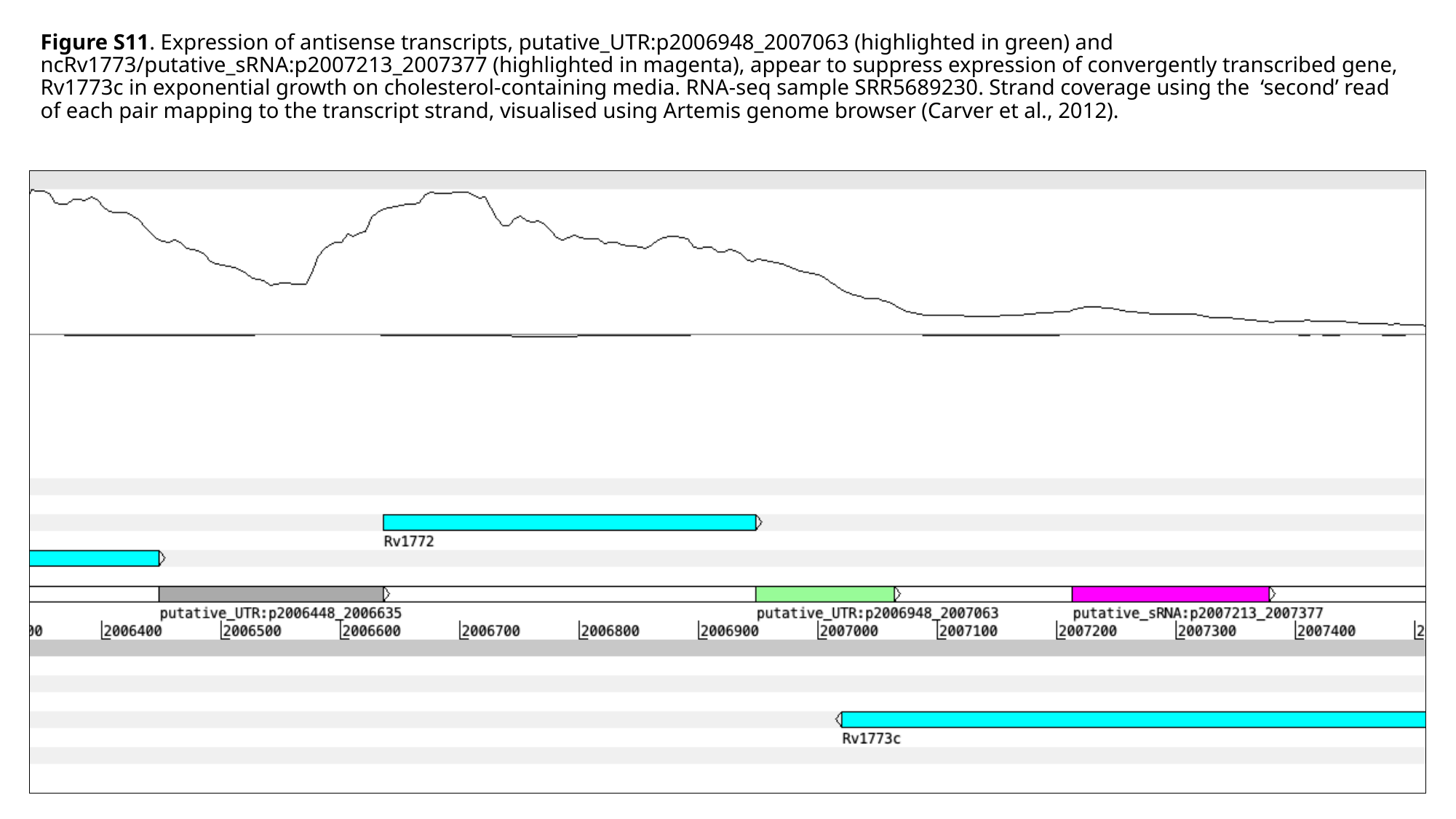

#### Figure S11. Expression of antisense transcripts, putative_UTR:p2006948_2007063 (highlighted in green) and ncRv1773/putative_sRNA:p2007213_2007377 (highlighted in magenta), appear to suppress expression of convergently transcribed gene, Rv1773c in exponential growth on cholesterol-containing media. RNA-seq sample SRR5689230. Strand coverage using the ‘second’ read of each pair mapping to the transcript strand, visualised using Artemis genome browser (Carver et al., 2012).

### Slide 13
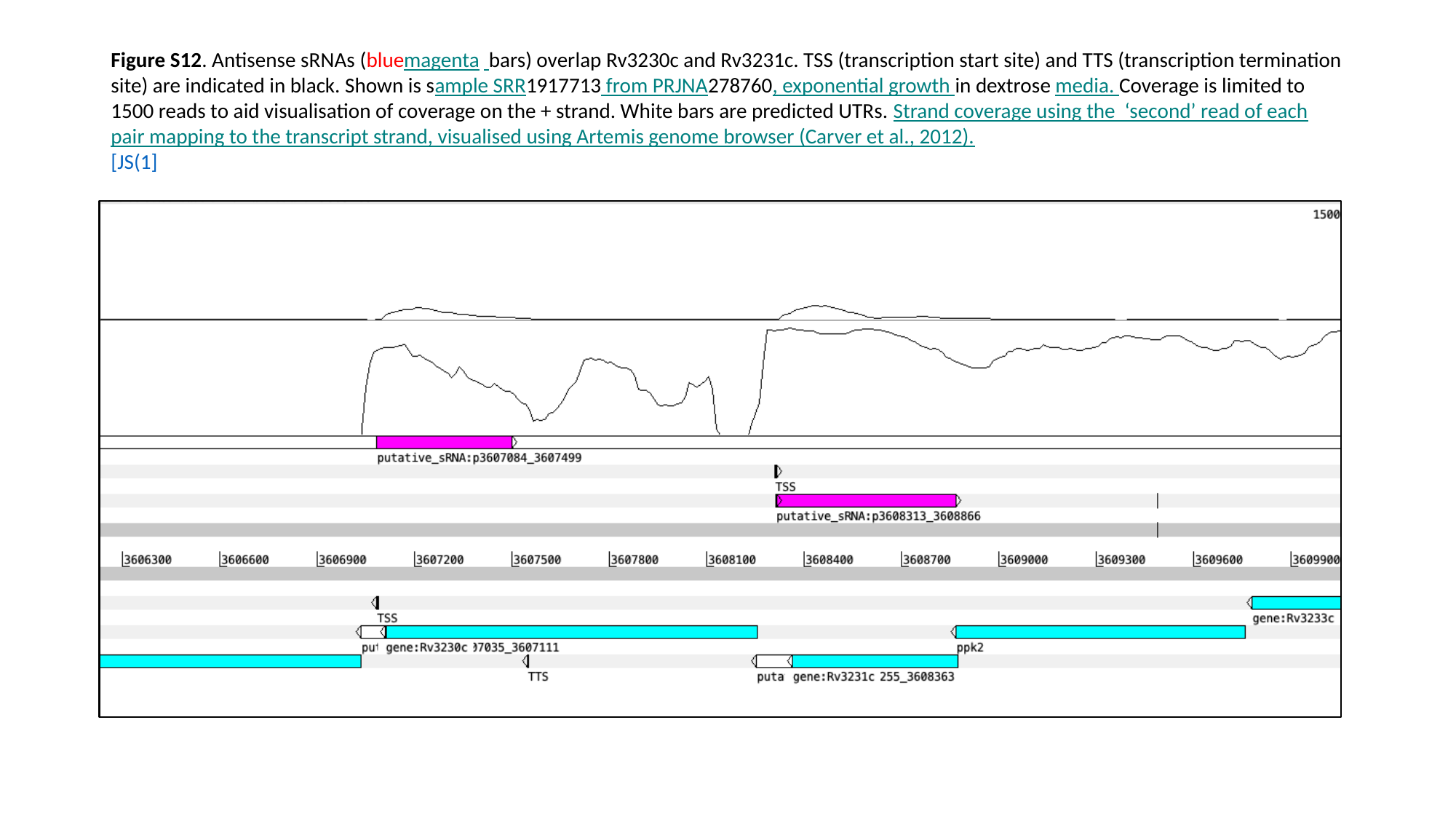

#### Figure S12. Antisense sRNAs (bluemagenta bars) overlap Rv3230c and Rv3231c. TSS (transcription start site) and TTS (transcription termination site) are indicated in black. Shown is sample SRR1917713 from PRJNA278760, exponential growth in dextrose media. Coverage is limited to 1500 reads to aid visualisation of coverage on the + strand. White bars are predicted UTRs. Strand coverage using the ‘second’ read of each pair mapping to the transcript strand, visualised using Artemis genome browser (Carver et al., 2012). [JS(1]

### Slide 14
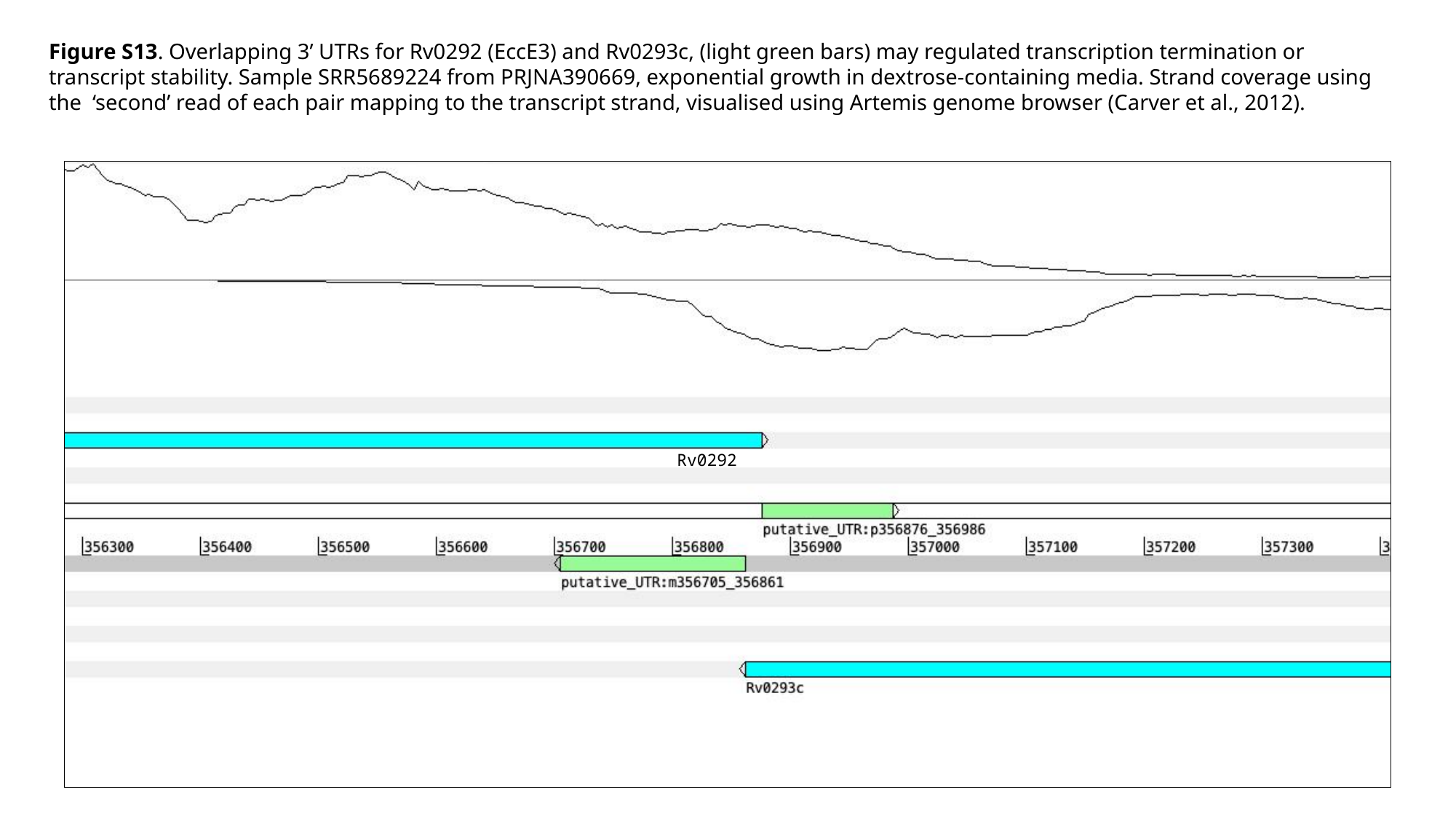

Figure S13. Overlapping 3’ UTRs for Rv0292 (EccE3) and Rv0293c, (light green bars) may regulated transcription termination or transcript stability. Sample SRR5689224 from PRJNA390669, exponential growth in dextrose-containing media. Strand coverage using the ‘second’ read of each pair mapping to the transcript strand, visualised using Artemis genome browser (Carver et al., 2012).
Rv0292
Rv0292
